# CDK12/13 inhibitor, CTX-439, suppresses tumor growth and potentiates BCL-2 family blockade

**DOI:** 10.64898/2026.02.20.706902

**Authors:** Hiroko Yamakawa, Raghda Khatab, Akio Mizutani, Shunsuke Ebara, Takaharu Hirayama, Sunao Tanaka, Yusuke Tarumoto, Midori Sugiyama, Seiichi Sugino, Mariko Takano, Hidekazu Tokuhara, Toshio Tanaka, Hiroshi Banno, Yasuyoshi Arikawa, Yukiko Fukui, Yuri Fujimoto, Shafiqul Islam, Masakazu Toi, Kosuke Kawaguchi, Daisuke Morishita, Kosuke Yusa

## Abstract

CDK12 and CDK13 (CDK12/13) regulate transcription by phosphorylating Serine 2 (S2) of the C-terminal domain of RNA polymerase II and have been proposed as therapeutic targets in cancer. Here we report the development of CTX-439, a novel, orally bioavailable, ATP-competitive small-molecule CDK12/13 inhibitor. CTX-439 specifically inhibits S2 phosphorylation and downregulates many genes including those involved in DNA damage repair, thereby exerting a profound anti-cancer effect *in vitro* and *in vivo* including breast cancer PDX models. A CRISPR activation screen identified *BCL-2* and *BCL-xL*, anti-apoptotic BCL-2 family members, as genes that when upregulated confer resistance to CTX-439. Simultaneous inhibition of BCL-2/BCL-xL and CDK12/13 rapidly induced apoptosis and significantly suppressed xenograft tumor growth. Mechanistically, CTX-439 downregulates MCL1 protein levels through transcriptional readthrough, shifting cell survival dependency to BCL-2 and BCL-xL. Our study provides novel insights into the anti-tumor effect of CDK12/13 inhibition and proposes a new combination therapy strategy with anti-apoptotic BCL-2 family inhibitors, which may improve therapeutic outcomes in cancer treatment.

## Introduction

Cyclin-dependent kinases (CDKs) are composed of 2 major classes, each playing a critical role in either cell-cycle regulation or transcription. Cell-cycle related CDKs, such as CDK1, CDK2 and CDK4/6, were explored as molecular targets in cancer therapy earlier. Inhibitors targeting CDK4/6 showed clinical significance and have been approved for the treatment of estrogen receptor (ER)-positive HER2-negative advanced or metastatic breast cancers in combination with an aromatase inhibitor (Beaver *et al*, 2015; Turner *et al*, 2018; Shah *et al*, 2018; Royce *et al*, 2022). The CDK4/6 inhibitors exhibited significant anti-tumor effects in other types of cancers and may thus be applicable in a wider range of tumors (Xue *et al*, 2019a, 2019b). Transcriptional CDKs include CDK7, CDK9, CDK11, CDK12, CDK13, and others, and inhibitors targeting CDK7 or CDK9 are currently under active clinical evaluation. Of these, the CDK7 inhibitors SY-5609 and samuraciclib, and the CDK9 inhibitor GFH009 have recently been granted orphan drug and/or fast track designation by the FDA (Zhou *et al*, 2023; Marineau *et al*, 2022; Patel *et al*, 2018). These inhibitors target CDKs that are involved in transcription by phosphorylating different sites of the carboxy-terminal domain (CTD) of RNA polymerase II (Pol II); CDK7 phosphorylates Serine 5 (S5) and Serine 7 (S7), which initiates and allows transcription to the proximal pausing sites. Following this, CDK9 phosphorylates S2 and enhances productive elongation beyond the proximal pausing site (Chen *et al*, 2018). CDK12 and CDK13 (CDK12/13) are important to maintain S2 phosphorylation for transcriptional elongation of particularly long genes (Harlen & Churchman, 2017). While pharmacological inhibition of CDK9 results in complete transcriptional shutdown, CDK12/13 inhibition affects transcription of long genes and results in premature termination due to intronic polyadenylation (Dubbury *et al*, 2018; Krajewska *et al*, 2019). DNA damage repair (DDR) genes are particularly enriched in long genes and carry intronic polyadenylation sites more frequently than average, which potentially makes them more susceptible to CDK12/13 inhibition (Blazek *et al*, 2011; Dubbury *et al*, 2018). A small subset of cancers (∼1%) in various cancer types carry CDK12 loss-of-function mutations with metastatic castration-resistant prostate cancer carrying more frequent mutations (∼6.9 %) (Robinson *et al*, 2015, 2017). Such tumors show characteristic Mb-sized tandem duplication phenotypes, further indicating the involvement of CDK12 in the DDR pathway (Sokol *et al*, 2019). These unique features of CDK12/13 inhibition have been exploited for cancer treatment; in addition to anti-cancer activity against various cancer types, it has been shown that CDK12/13 inhibition can re-sensitize cells with acquired resistance to PARP1 inhibition (Johnson *et al*, 2016) and that CDK12-deficient tumors may be sensitive to immunotherapy due to higher neoantigen burden (Wu *et al*, 2018), representing broad applicability of CDK12/13 inhibition.

Several compounds targeting CDK12/13 have been developed: THZ531 is a covalent inhibitor (Zhang *et al*, 2016) and SR4835 inhibits CDK12/13 through a molecular glue activity that induces proteasomal degradation of Cyclin K (Quereda *et al*, 2019; Houles *et al*, 2023). These compounds showed anti-tumor activity by downregulating DDR genes and inducing apoptosis. Together with a growing number of CDK12/13 or partner Cyclin K targeting agents, various studies have proposed targeted therapies against CDK12/13 in a wide range of tumors (Iniguez *et al*, 2018; Quereda *et al*, 2019).

Here we describe the novel investigational new drug (IND)-ready CDK12/13 inhibitor, CTX-439, with an IC_50_ of 3.1 nM in cell-free assays. CTX-439 treatment specifically inhibited S2 phosphorylation of Pol II CTD and downregulated a large number of genes, including DDR genes, which induced marked anti-tumor effects *in vitro* as well as *in vivo*. CTX-439 treatment acutely downregulated anti-apoptosis protein, MCL1, which when combined with BCL-2 and BCL-xL inhibition induced profound apoptosis. We propose monotherapy with CTX-439 as well as combination treatment with anti-apoptotic BCL-2 family inhibitors for a more effective CDK12/13-based therapy.

## Results

### Novel CDK12/13 inhibitor, CTX-439, exhibits anti-tumor activity *in vitro* and *in vivo*

In order to establish a CDK12/13-targeted cancer therapy, we developed a novel orally bioavailable CDK12/13 inhibitor, CTX-439 (Fig. 1A and Supplementary Fig. 1), which showed a high binding affinity towards CDK12 (Kd=0.38nM; Fig. 1B and Supplementary Data 1) and exhibited the half maximal inhibitory concentration (IC_50_) of 3.1 and 9.2 nM towards CDK12/Cyclin K and CDK13/Cyclin K, respectively, in cell-free assays (Fig. 1C and Supplementary Table 1). CTX-439 possesses a highly specific inhibitory activity towards CDK12/13 compared to other members of the CDK family proteins, showing > 550-fold higher binding affinity and 20-fold higher inhibitory activity than the closest family member, CDK9 (Fig. 1B-C). Among a 468 kinase panel, CTX-439 showed cross-reactivity against ASK1, DYRK1A and TAOK1/3 (Fig. 1B and Supplementary Table 1). CTX-439 also showed an inhibitory activity superior to two existing inhibitors (Fig. 1C)

**Fig. 1.**
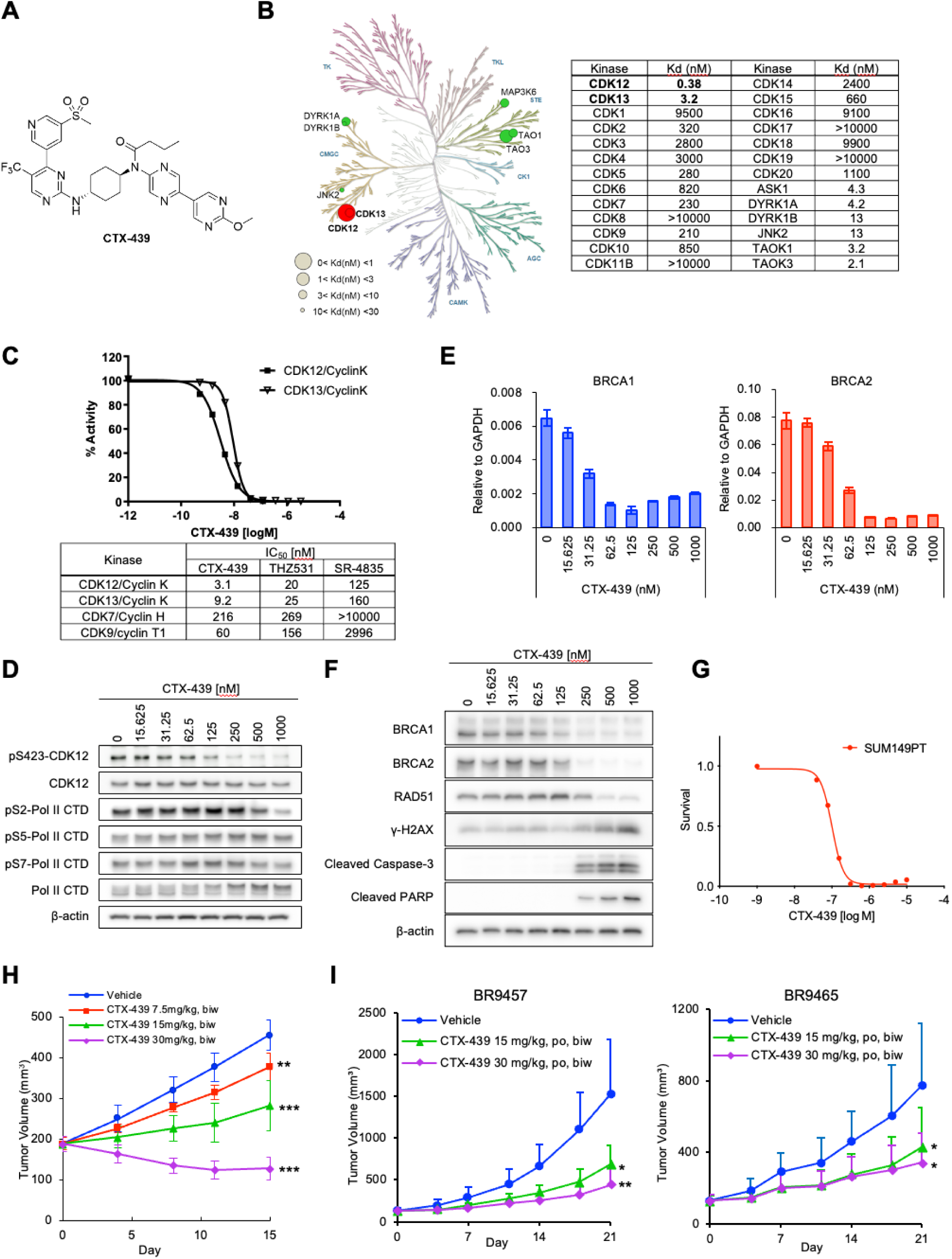
Generation of a novel CDK12/13 inhibitor, CTX-439 and its anti-tumor potential. (**A**) Chemical structure of CTX-439. (**B**) Kinome tree illustrating the selectivity profile of CTX-439. Binding affinity of CTX-439 was screened by scanMAX kinase panel and the Kd values for CDKs and the other top 6 kinases were evaluated. For CASK, DCAMKL3, DYRK2, MAP3K15, JNK3, JNK1, RSK4, ERK8, ERN1, RSK3 and CLK1, CTX-439 showed weak inhibitory effects of 50-65% at 30 nM, but it is not included in the kinome tree. (**C**) Inhibitory activities of CTX-439, THZ531 and SR4835 on transcription-related CDK kinases. (**D**) Western blot analysis of CDK12 and Pol II CTD phosphorylation in SUM149PT cells treated with the indicated concentration of CTX-439 for 6 h. (**E**) RT-qPCR analyses of *BRCA1* and *BRCA2* in SUM149PT cells treated with CTX-439 for 6h. (**F**) Western blot analysis of DDR proteins and apoptosis markers in SUM149PT cells treated with CTX-439 for 24h. (**G**) Drug response assay in SUM149PT cells treated with CTX-439 for 72h. (**H**) *in vivo* anti-tumor efficacy of CTX-439 in mice bearing SUM149PT xenografts treated with the indicated dose twice weekly (biw). (**I**) *in vivo* anti-tumor efficacy of CTX-439 in TNBC PDX model mice treated with the indicated dose. * *p*<0.05, ** *p*<0.01 ** *p*<0.001 by Dunnett’s test. Data are shown as mean ± SD (n=3, **E**; n=6, **H** and **I**).

CDK12/13 primarily regulate phosphorylation of S2 (p-S2) in the CTD of Pol II and play a critical role in transcriptional elongation and splicing. To further investigate the inhibitory activity of CTX-439, we analyzed the status of key phosphorylation sites within the CTD in breast cancer cell line, SUM149PT, treated with CTX-439. As shown in Figure 1D, CTX-439 downregulated p-S2, but neither phosphorylation of S5 (p-S5) nor phosphorylation of S7 (p-S7), supporting its specificity. Serine 423 (S423) of CDK12 is a self-phosphorylation target (Yamakawa *et al*, 2024) and CTX-439 also reduced its phosphorylation level in a dose-dependent manner (Fig. 1D). It has been shown in genetic and pharmacological studies that CDK12 plays a role in the expression of DDR genes (Dubbury *et al*, 2018).

Consistent with this notion, CTX-439 treatment downregulated *BRCA1/2* expression (Fig. 1E), which was also confirmed at the protein level (Fig. 1F). Rad51 downregulation was also detected (Fig. 1F). This DDR gene downregulation was accompanied with upregulation of double-strand break marker, γH2AX, and apoptosis marker, cleaved PARP (cPARP) and cleaved Caspase-3 (cCasp3) (Fig. 1F). We further assessed the effects on transcription by RNA-seq and found a marked downregulation with particular enrichment of the DDR pathway (Supplementary Fig. 2 and Supplementary Data 2). These results strongly suggest that CTX-439 possesses profound anti-tumor effects. Indeed, we observed an IC_50_ of 100.5 nM against SUM149PT in cell-based assay (Fig. 1G). This observation was further extended to multiple cell lines derived from breast and ovarian cancer, indicating a broad anti-proliferation effect by CTX-439 (Supplementary Table 2).

To investigate the *in vivo* efficacy of CTX-439, we treated nude mice bearing SUM149PT xenografts by oral gavage twice weekly for 2 weeks. We found that CTX-439 showed marked *in vivo* anti-tumor activity, accompanied by tumor regression at the highest dose tested (Fig. 1H). Investigation of the pharmacokinetics of CTX-439 in plasma and tumors showed rapid accumulation, reaching its highest concentration within 1 and 2 h post administration, respectively, and declined with a half-life of approximately 4 h (Supplementary Fig. 3A-B). *BRCA1/2* expression in tumors, measured by RT-qPCR, showed rapid downregulation, but as drug concentration was decreased, the expression of these downstream genes recovered, albeit at different kinetics (Supplementary Fig. 3C-D). At the highest dose, *BRCA1/2* expression was kept downregulated by > 50 % for approximately 16 h. Despite the clear drug engagement, mice treated with CTX-439 did not show significant weight loss at all doses tested (Supplementary Fig. 3E), suggesting that CDK12/13 inhibition by CTX-439 was well tolerated in mice. We further tested the anti-tumor efficacy of CTX-439 using breast cancer patient-derived xenograft (PDX) models (1 ER+, 1 ER+/HER2+ and 6 triple-negative breast cancer (TNBC) models; Supplementary Tables 3 and 4). CTX-439 treatment was most effective in suppressing TNBC tumors (Fig. 1I and Supplementary Fig. 4).

Taken together, we successfully developed the novel CDK12/13 inhibitor, CTX-439, with anti-tumor activity against a broad range of breast and ovarian cancer cell lines as well as pronounced *in vivo* anti-tumor activity.

### BCL-xL inhibition boosts the cytotoxic activity of CTX-439

Molecular target therapy often faces acquired resistance and it is critically important to identify biomarkers that predict treatment outcome for effective therapy. To identify genes that affect anti-tumor efficacy of CTX-439, we performed a CRISPR-activation screen in SUM149PT (Supplementary Data 3). We identified 69 genes, which confer resistance when upregulated (log fold change (LFC) > 0.5 in CTX-439-vs-CTRL, LFC < 0.5 in DMSO-vs-CTRL and positive p-value < 0.01 in CTX-439-vs-CTRL). The top hit was *ABCB1*, which encodes a member of ABC transporters. Interestingly, *CDK12* was a significant hit, but *CDK13* was not. Although these 2 kinases largely share functions, CDK12 seems to be more important to the survival of cells under CTX-439 treatments. Genes that caught our attention the most were those encoding anti-apoptosis proteins, *BCL-2* and *BCL-xL* (also known as *BCL2L1*), which ranked within the top 5 (Fig. 2A). Interestingly, *MCL1*, another anti-apoptosis gene, did not confer resistance. These results suggest that BCL-2/xL may limit anti-tumor efficacy of CTX-439 and that suppression of them may enhance the anti-tumor effect. We therefore tested dual inhibition with CTX-439 and AZD4320 (BCL-2/xL inhibitor) and found a significant synergistic effect in SUM149PT (Fig. 2B) and the ovarian cancer cell line OV90 (Supplementary Fig. 5A). THZ531, another CDK12/13 inhibitor, also showed synergistic effect with AZD4320 (Supplementary Fig. 5A). A-1331852, a BCL-xL-specific inhibitor, also synergized with CTX-439 (Supplementary Fig. 5B). These strong cell killing effects were induced within 8h, which reduced cells by ∼90% at the highest dose tested (Fig. 2B-D, Supplementary Fig. 5C-E). To analyze cell death kinetics, we performed a time-course cell count assay and found that significant cell death was detected as early as 4h (Fig. 2E and Supplementary Fig. 5F). The use of BCL-xL inhibitors and the rapid onset of cell death strongly suggest that the synergistic effect is mediated by acute induction of apoptosis. As expected, a significantly higher percentage of Annexin-V positive cells were detected in the combined treatment, while CTX-439 or AZD4320 alone were not effective in inducing apoptosis within this time range (Fig. 2F and Supplementary Fig. 5G).

**Fig. 2.**
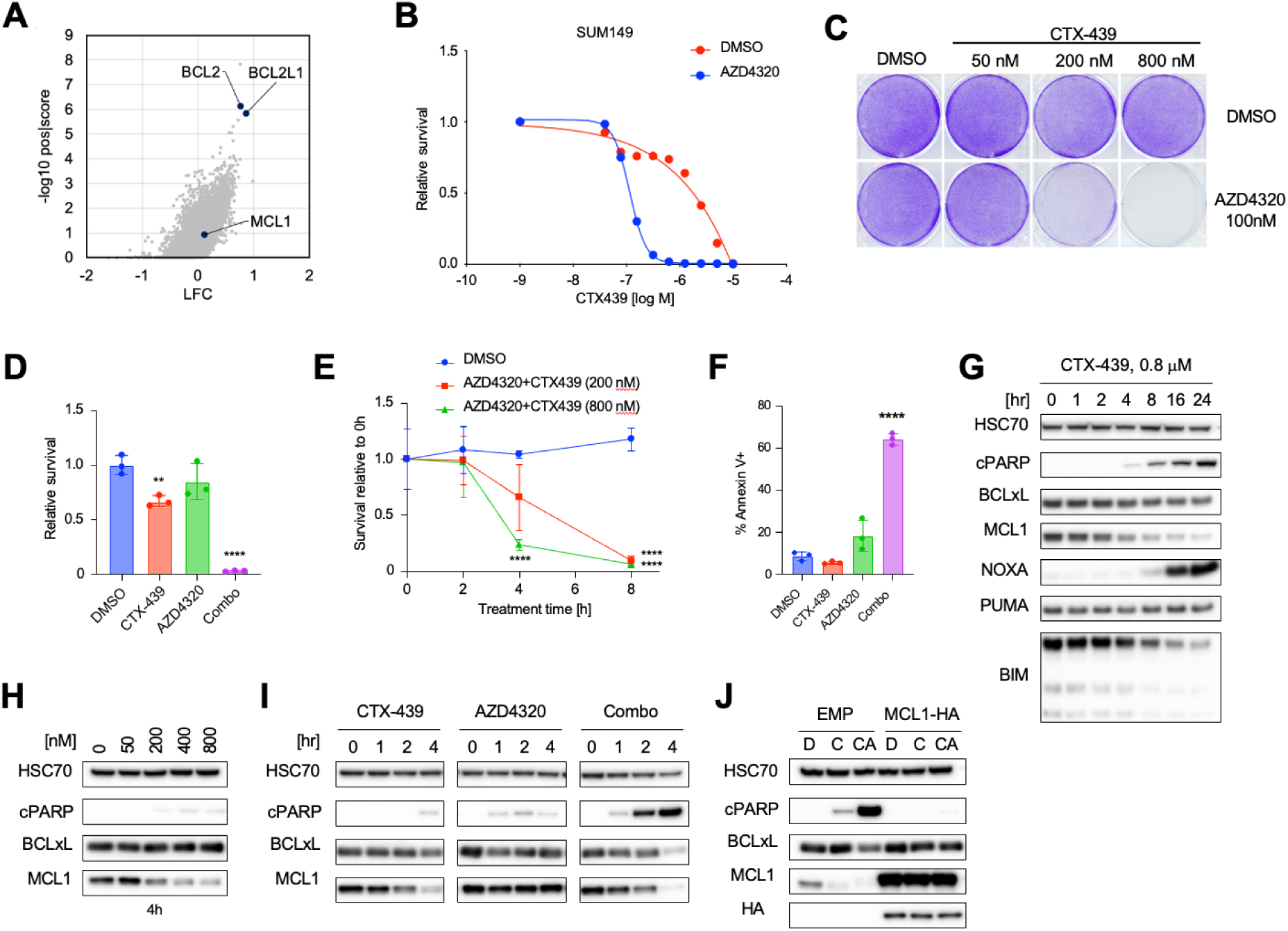
BCL-2/xL inhibition exhibited a profound synergistic effect with CDK12/13 inhibition. (**A**) Volcano plot showing the positive selection outcome of the CRISPR-activation screen using SUM149PT cells treated with CTX-439. Anti-apoptosis genes are highlighted. (**B**) Cell survival assay. SUM149PT cells were treated with serially diluted CTX-439 together with DMSO or AZD4320 for 8h, and then further incubated for 64h without treatment. (**C**) Crystal violet staining of cells treated in the indicated drug combination for 8h followed by 40h incubation without treatment. (**D-E**) Relative cell counts 8h post treatment (**D**) or at the indicated time point (**E**). (**F**) Annexin V staining. Cells were treated with either CTX-439 or AZD4320, or in combination for 6h. **G**. Western blot analysis of BCL-2 family proteins. Cells were treated with CTX-439 for the indicated time. (**H-I**) Western blot analysis of anti-apoptosis proteins and apoptosis marker, cleaved PARP (cPARP). Cells were treated at the indicated concentration of CTX-439 for 4h (**H**) or for the indicated time (**I**). (**J**) Western blot analysis of MCL1-overexpressing cells treated with DMSO (D), CTX-439 alone (C) or in combination with AZD4320 (CA). Exogenous MCL1 was tagged with HA. Unless otherwise specified, CTX-439 at 800 nM and AZD4320 at 100 nM were used. Data are shown as mean ± SD (n=3, **D-F**). * *p*<0.05, ** *p*<0.01, *** *p*<0.001 by Student’s *t-*test.

In order to reveal the mechanism of the synergy between CDK12/13 and BCL-2/xL inhibition, we profiled the expression kinetics of major BCL-2 family proteins up to 24 h post CDK12/13 inhibition. When treated with CTX-439 alone, both SUM149PT and OV90 showed downregulation of the anti-apoptotic protein, MCL1, starting at as early as 2 h (Fig. 2G and Supplementary Fig. 5H). Beyond 2 h, MCL1 was downregulated further to a lower level, although it slightly recovered in OV90 from 16 h onward. THZ531 treatment in OV90 also downregulated MCL1 (Supplementary Fig. 5I) and was dose-dependent (Fig. 2H and Supplementary Fig. 5J). Among pro-apoptosis proteins tested, BIM was downregulated in both cell lines from 4 h onward and NOXA was upregulated at 16 h, albeit at different levels between the 2 cell lines. Consistent with Annexin-V staining, cPARP expression was very weak when cells were treated with CTX-439 or AZD4320 alone, even though MCL1 was significantly downregulated by CTX-439 treatment at 4 h (Fig. 2I and Supplementary Fig.5K). In contrast, when the two inhibitors were combined, cPARP was rapidly and significantly upregulated, which negatively correlated with MCL1 downregulation (Fig. 2I and Supplementary Fig. 5K). To confirm whether MCL1 downregulation via CDK12/13 inhibition is responsible for rapid apoptosis induction in combination with BCL-xL inhibition, we overexpressed MCL1 and tested apoptosis induction. In both cell lines, exogenous expression of MCL1 completely suppressed cPARP expression (Fig. 2J and Supplementary Fig. 5L). We also confirmed that the MCL1 inhibitor, AZD5991, did not affect the anti-proliferation effect of CTX-439 (Supplementary Fig. 5M). Rapid apoptosis induction by suppression of BCL-2/xL and MCL1 is likely through the mitochondrial pathway, in which BAK and BAX mediate cytochrome c release and activate downstream caspases. We therefore tested if combined treatment could induce apoptosis in *BAK/BAX* double knockout cells and found that cell death and apoptosis induction was significantly suppressed (Supplementary Fig. 6). Taken together, we found that CDK12/13 inhibition induces rapid downregulation of MCL1 which can be exploited for combination therapy with BCL-xL inhibition. Indeed, MCL1 downregulation is observed in a wide range of cancer cell lines, suggesting broad applicability (Supplementary Fig. 7).

### CDK12/13 inhibition causes aberrant *MCL1* transcription

Given that CDK12/13 are involved in Pol II regulation by phosphorylating S2 of the CTD, rapid MCL1 downregulation at the protein level is most likely caused by dysregulated transcription. However, RNA-seq analysis using total RNA did not show a significant change in *MCL1* at the transcript level (Supplementary Fig. 2). We therefore performed RNA-seq analysis using fractionated RNA, namely nuclear and cytoplasmic RNAs, extracted from DMSO or CTX-439 treated cells. For nuclear RNA, we generated RNA-seq libraries using the polyA+ fraction as well as rRNA-depleted RNAs. Shown in Figure 3A are RNA-seq tracks of the *MCL1* locus. The most noticeable abnormality was reads accumulated beyond the 3’ end of the *MCL1* gene. In the analysis using rRNA-depleted RNAs, reads farther than 20-kb downstream, which completely cover the neighboring gene *ADAMTSL4*, were detected in the same orientation as *MCL1* transcription. These reads were therefore most likely to have originated from readthrough transcription of *MCL1*. Analysis of polyA+ nuclear RNAs also detected readthrough, albeit at a shorter length, suggesting that a fraction of the mature mRNA carry 3’ extension of the untranslated region (UTR). Analysis of cytoplasmic RNA however did not show as many readthrough reads as nuclear RNA. Intriguingly, the *MCL1* expression levels calculated based on exonic reads revealed an opposite change between the subcellular fractions: upregulation in the nucleus, but downregulation in the cytoplasm (Fig. 3A. Note that the *y*-axis ranges are not equal). In addition to the 3’ end abnormality, we noticed that in rRNA-depleted nuclear RNA, intronic reads relative to the adjacent exons, especially in Intron 1, were higher in CTX-439 treated cells (Fig. 3B), indicating splicing abnormality.

**Fig. 3.**
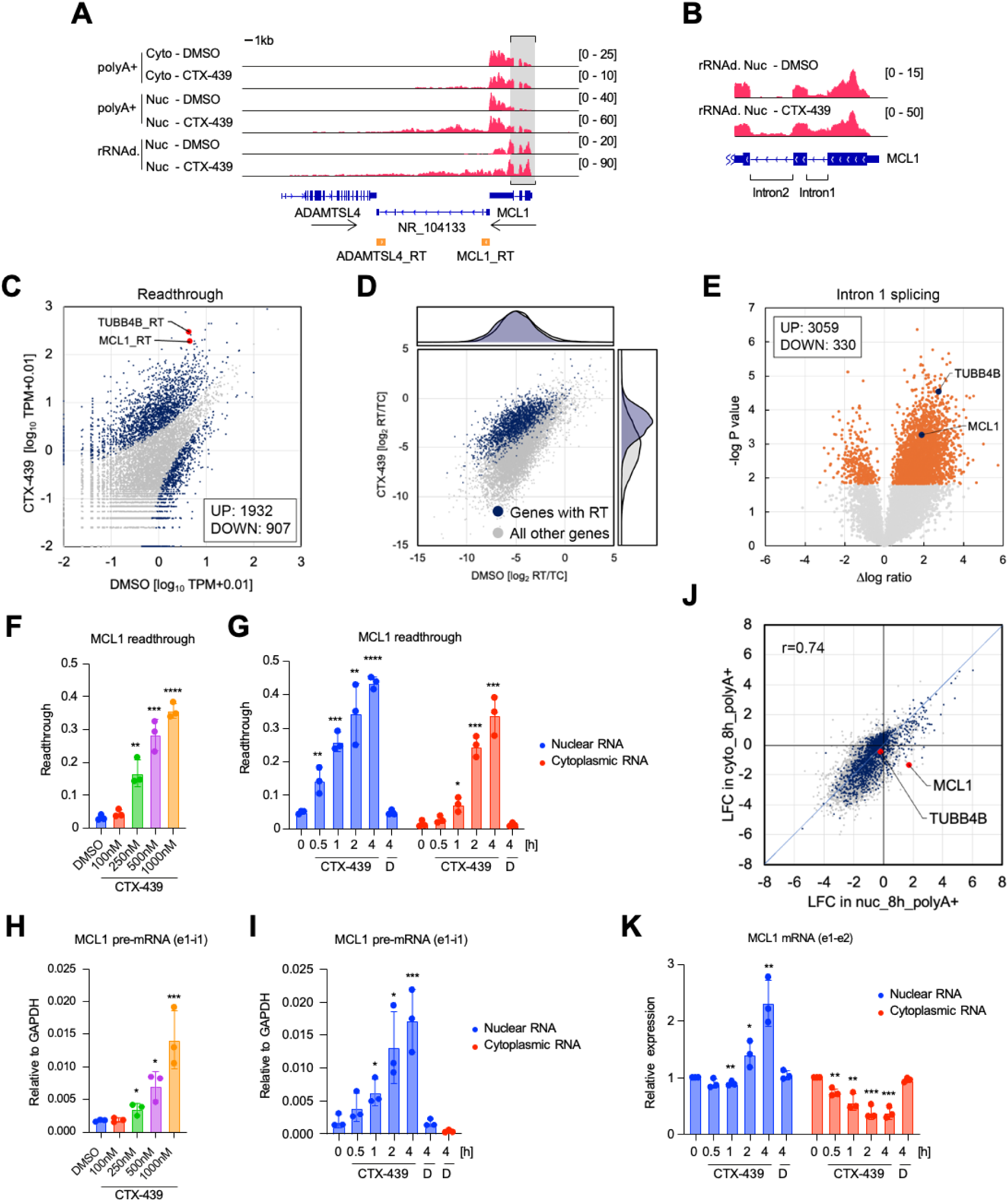
CTX-439 treatment induced transcriptional readthrough and splicing defects. (**A**) RNA-seq track showing *MCL1* expression from the indicated drug treatment for 8h and RNA preparation. Cyto, cytoplasmic RNA; Nuc, nuclear RNA; polyA+, polyA+ enriched RNA, rRNAd, rRNA-depleted RNA. Note that the *y* axis hights are ajusted as indicated for easier comparisons. Red peaks indicate reads from negative strand. Positive-strand track is omitted as no reads were detected in this region. *AMADTSL4* and *MCL1* are coded on positive and negative strand, respectively. 1-kb regions in which aligned reads are counted as readthrough are indicated in orange with the _RT suffix. (**B**) Magnified track of the region indicated in (**A**). (**C**) Scatter plot comparing readthrough counts in cells treated with DMSO or CTX-439. Significant genes are shown in blue. (**D**) Scatter plot showing ratios of readthrough counts to corresponding coding transcript counts. Genes with significantly overexpressed readthrough are highlighted in blue, indicating increased ratios compared to the non-significant genes. (**E**) Volcano plot showing DIntron1-to-Exon1 ratios (CTX-439 - DMSO) compared to statistical significance. Significantly altered genes (*p*<0.05) are highlighted in orange. Statistical analysis was performed with Welch’s test. (**F-G**) Dose- (**F**) and time-series (**G**) RT-qPCR analysis of *MCL1* readthrough transcription. Cells were treated with DMSO (D) or CTX-439 for 4h at the indicated concentration (**F**) or at 1 μM for the indicated time (**G**). Nuclear and cytoplasmic RNA were then isolated. (**H**-**I**) Dose- (**H**) and time-series (**I**) RT-qPCR analysis of *MCL1* pre-mRNA expression. The Exon1-Intron1 junction was measured. (**J**) Scatter plot comparing expression LFC in the nucleus (*x*) and that in cytoplasm (*y*). Note that *MCL1* is upregulated in the nucleus but downregulated in the cytoplasm. (**K**) Time-series RT-qPCR analysis of the *MCL1* transcript level in nuclear and cytoplasmic RNAs. Data are shown as mean ± SD (n=3, **F-I** and **K**). * *p*<0.05, ** *p*<0.01, *** *p*<0.001 by Student’s *t-*test.

Global analysis of readthrough transcription (summarized in Supplementary Fig. 8A-D) revealed that, upon CTX-439 treatment, approximately 2,000 genes significantly increased read counts mapped downstream of their last exon, while 900 genes decreased counts (Fig. 3C). Log2 ratios of readthrough counts to exonic counts of corresponding genes were similar between the up-regulated and non-significant genes in DMSO-treated cells; however, when cells were treated with CTX-439, this ratio became higher in the up-regulated genes (Fig. 3D), suggesting that the increase in readthrough reads was not simply due to transcriptional upregulation, but rather defects in the 3’ end processing. We additionally performed global analysis on splicing by focusing on intron 1 (summarized in Supplementary Fig. 9). This analysis revealed that more than 3,000 genes showed significantly higher read counts mapped in Intron 1 upon CTX-439 treatment (Fig. 3E), indicating wide-spread splicing defects, although there were a subset of genes exhibiting improved splicing efficiency (Supplementary Fig. 9C).

qPCR analysis revealed that readthrough transcription in *MCL1* and another high readthrough gene, *TUBB4B,* was dose-dependent (Fig. 3F and Supplementary Fig. 8E-F) and the acute response was detected as early as 30 min post treatment, followed by further increase until at least 4 h, in the nuclear fraction (Fig. 3G and Supplementary Fig. 8G). In the cytoplasm, the increase of *MCL1* 3’ readthrough was detected in a time-dependent manner with an initial delay compared to nuclear RNA, but the increase was much weaker in cytoplasmic *TUBB4B* transcript (Fig. 3G and Supplementary Fig. 8G). *MCL1* splicing defects also showed dose- and time-dependent increase (Fig. 3H-I). These increased readthrough and splicing defects in *MCL1* transcription were detected in a wide range of cancer cell lines when treated with CTX-439 (Supplementary Fig. 10).

As mentioned previously, the change of the *MCL1* transcript level upon CTX-439 treatment was opposite between the nucleus and the cytoplasm (Fig. 3A). Globally, however, transcriptional changes showed a general agreement between nuclear and cytoplasmic RNAs (Fig. 3J; Pearson r=0.74). Therefore, *MCL1* seemed to be a rare exception. qPCR analysis of spliced transcripts (e1-e2 junction) revealed that in the nucleus, *MCL1* transcripts showed a slight decrease for the first 1 h, followed by a significant increase from 2 h onward, but the cytoplasmic level continuously decreased upon treatment (Fig. 3K). In summary, CTX-439 treatment induces abnormalities in splicing and 3’ end processing, and causes a unique expression pattern of *MCL1*, resulting in rapid cytoplasmic downregulation at both the mRNA and protein levels.

### Defective 3’ end processing causes cytoplasmic downregulation of *MCL1* transcripts

Next we sought to further investigate the discrepancy of the *MCL1* transcript level between the nucleus and the cytoplasm. Focusing on the acute phase (first 1 h) post treatment, spliced mRNA (e1-e2 junction) mostly kept its expression level in the nucleus, while it decreased in the cytoplasm (Fig. 3K). During this period, the rapid increase of readthrough transcripts was detected (Fig. 3G). Although splicing was certainly affected, these observations suggest that the defect in 3’ end processing, rather than splicing, makes a significant impact on MCL1 expression. In addition, it is known that the 3’ UTR of the *MCL1* mRNA contains destabilization signals that increase mRNA turnover rate, thereby keeping the *MCL1* mRNA and protein levels low (Senichkin *et al*, 2020). These may combinatorially affect the cytoplasmic mRNA level and consequently the protein level.

To investigate the involvement of 3’ end processing and the 3’ UTR, we inserted a polyadenylation signal sequence (bovine growth hormone pA, bpA) widely used in synthetic gene cassettes, together with a selection marker, immediately before the termination codon of *MCL1* at the endogenous locus (Fig. 4A). In both SUM149PT and OV90, bpA insertion significantly increased the basal level of MCL1 protein (Fig. 4B). Upon CTX-439 treatment, while *MCL1* wild-type cells decreased the MCL1 protein level, cells with bpA insertion maintained or even slightly increased MCL1 protein level (Fig. 4B). Since MCL1 downregulation was no longer induced by CTX-439 treatment in the bpA-inserted cells, apoptosis induction (cPARP) by combination treatment was significantly suppressed in SUM149PT and not even detected in OV90 (Fig. 4B). Consistent with the protein level, the basal expression level of *MCL1* mRNA in the cytoplasm was upregulated by bpA insertion, most likely due to a disconnection from the destabilization signals (Fig. 4C-D). Upon CTX-439 treatment, the cytoplasmic *MCL1* mRNA level was further increased in the bpA-inserted cells (Fig. 4C-D), while reproducibly downregulated in wild-type cells. Therefore, bpA insertion gave rise to a striking difference in cytoplasmic mRNA expression.

**Fig. 4.**
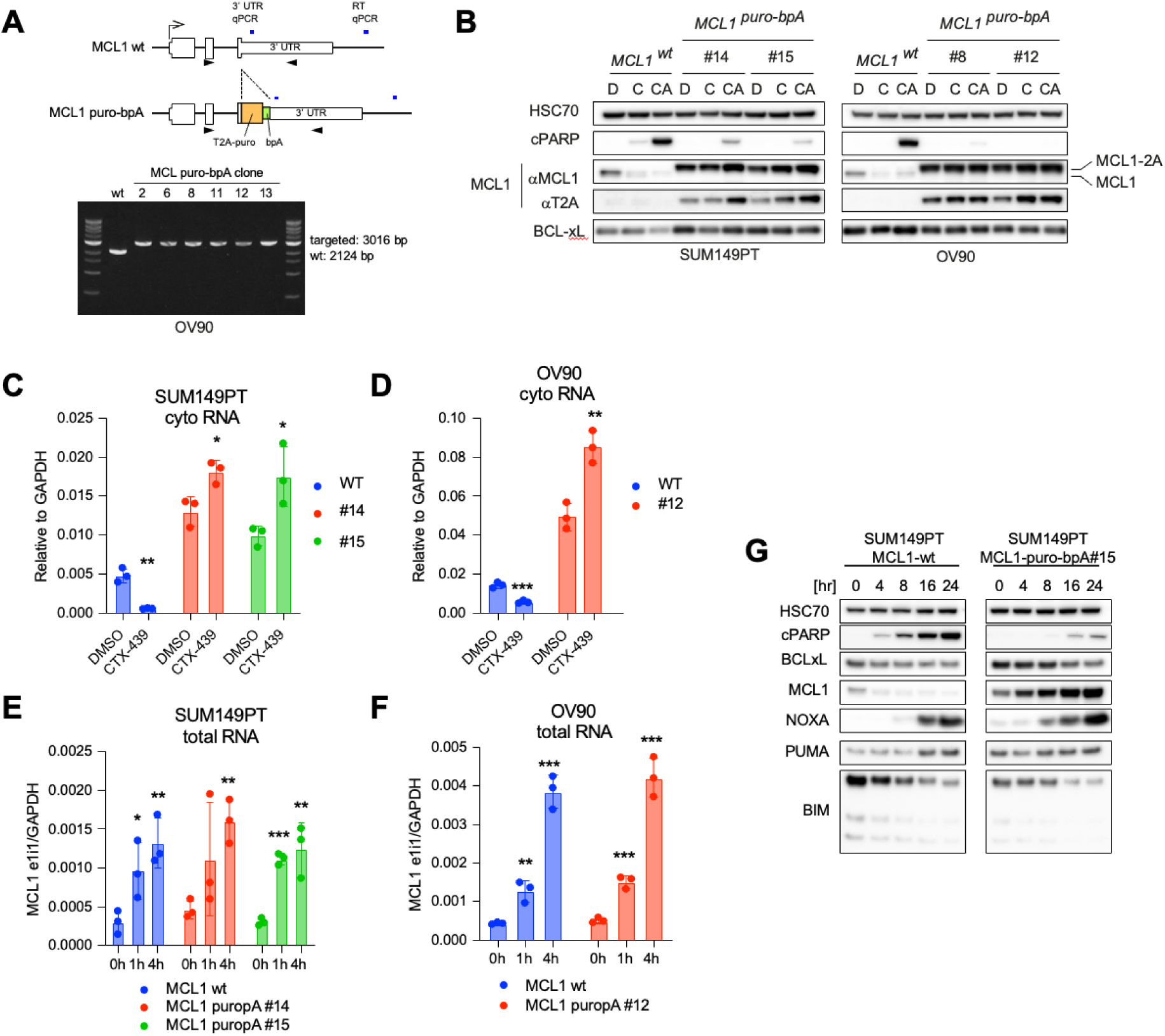
*MCL1* 3’ UTR causes CTX-439-induced cytoplasmic downregulation. (**A**) Schematic of the endogenous *MCL1* locus modification. The T2A-puro-bpA cassette was replaced with the termination codon. Bottom photo shows homozygous knock-in. Primers (arrowheads) were used. (**B**) Western blot analysis of cPARP, MCL1, T2A (targeted MCL1) and BCL-xL in SUM149PT (left) and OV90 (right) clones treated with DMSO (D), CTX-439 (C) or in combination of CTX-439 and AZD4320 (CA) for 4h. (**C**-**D**) RT-qPCR analysis of the cytoplasmic *MCL1* transcript level. Cells were treated with CTX-439 for 4h and cytoplasmic RNA was purified. (**E**-**F)** RT-qPCR analysis of intron 1 splicing. Cells were treated with the CTX-439 for the indicated time and total RNA were extracted. Pre-mRNA (junction of exon 1 and intron 1) was analyzed. (**G**) Western blot analysis of SUM149PT wildtype and clone 15. Lack of MCL1 down-regulation suppressed apoptosis induction. Data are shown as mean ± SD (n=3, **C-F**). * *p*<0.05, ** *p*<0.01, *** *p*<0.001 by Student’s *t-*test.

To assess whether bpA insertion affects transcriptional activity and/or splicing, we quantified pre-mRNA levels by measuring the e1-i1 junction using total RNA. qPCR analysis showed a time-dependent increase with no significant difference between wild-type and the bpA-inserted cells (Fig. 4E-F), indicating that bpA insertion affects neither transcription nor splicing. At the protein level, a longer treatment for up to 24 h gradually upregulated MCL1 in the bpA-inserted cells (Fig. 4G). Other BCL-2-family proteins tested showed no obvious differences except minor expression level changes, probably due to compensatory reactions from MCL1 upregulation (Fig. 4G). With this sustained expression of MCL1, apoptosis induction by CTX-439 monotherapy was nearly completely abolished (Fig. 4G), suggesting that MCL1 downregulation by inefficient 3’ end processing, to some extent, mediates the anti-tumor effect of CTX-439.

Taken together, these results indicate that the defects in 3’ end processing and the destabilization signal in the 3’ UTR cause cytoplasmic downregulation of *MCL1* mRNA in spite of transcriptional upregulation in the nucleus, leaving *MCL1* to a unique position where transcript levels alter to opposite directions between the nucleus and the cytoplasm.

### CTX-439 causes gene-length dependent abnormal transcription

CDK12/13 inhibition in principle gives a negative impact on transcription, especially to long genes (Dubbury *et al*, 2018). However, like *MCL1* shown above, there are genes whose expression is significantly upregulated under the CDK12/13 inhibition. Recent studies on CDK12/13 functions showed that transcriptional abnormalities depend on gene body size (Krajewska *et al*, 2019). To confirm whether such size-dependent effects can be seen in CTX-439 treated cells, we analyzed the RNA-seq data generated with the rRNA-depleted nuclear RNAs in further detail. We calculated the ratio of the mean per-base read depth of each intron or readthrough (here, defined by a 3-kb region immediately after the last exon) to the mean per-base read depth of the entire genome, and then plotted the calculated ratios along the distance from TSS to the mid-point of each intron/readthrough (Fig. 5A). This analysis showed that mean ratios were consistent across a wide range of distances from TSS for both intron and readthrough in control cells (Fig. 5A, top). In CTX-439 treated cells, however, mean ratios decreased at a farther distance, starting from several kb (Fig. 5A, middle). Subtraction between control and CTX-439 treated cells revealed a clear pattern that both intron and readthrough reads within an approximately 10-kb region from TSS were higher but decreased beyond 10 kb, in CTX-439 treated cells (Fig. 5A, bottom). We then grouped genes into 3 bins by gene body size and generated metagene plots with RPM (read per million; Fig. 5B). For genes with < 20 kb in size, intronic and readthrough read counts across the entire gene body and regions beyond TES were significantly higher in CTX-439 treated cells than control. In contrast, for genes with 20-80 kb and > 80 kb in size, read counts immediately after TSS were higher in CTX-439 treated cells, but decreased towards and beyond TES. As this analysis used read counts of introns, defects in splicing of proximal introns might have had a major impact on the increase observed immediately after TSS in any gene size. However, the data suggest higher transcriptional activity in 5’ regions, most likely up to 20 kb or more. If the 3’ end of a gene is located within this range (i.e. short genes), high incidents of readthrough transcription occur due to inefficient 3’ end processing. Consistent with this notion, genes with upregulated readthrough (median gene size of 6.3 kb) were significantly shorter than those with no-change (28.8 kb) and downregulated (73.4 kb) readthrough (Fig. 5C). As transcriptional upregulation near TSS is common to all gene classes, splicing inefficiency, especially of intron 1, did not correlate with gene body size (Fig. 5D).

Taken together, CTX-439 mediated CDK12/13 inhibition causes length-dependent transcriptional abnormalities, the defect in transcriptional elongation of particularly long genes and inefficient 3’ end processing of short genes, all of which align with those induced by THZ531(Krajewska *et al*, 2019).

### Transcriptional phenotypes are caused by genuine on-target effects of CTX-439

Although CTX-439 is highly specific to CDK12/13 (Fig. 1) and the experimental results support that the observed phenotypes were mediated by CDK12/13 inhibition, off-target activity of inhibitors is a concern for safety in clinical application. Also, targeted therapies with kinase inhibitors often lose their effectiveness due to gatekeeper mutations, the mechanism by which compounds are no longer able to bind to and effectively inhibit target proteins. To address these issues, we sought to identify CDK12 variants that confer resistance to CTX-439 without affecting CDK12 kinase activity. We designed multiple gRNAs targeting the CDK12 kinase domain and expressed them in Cas9-expressing MDA-MB-436 cells. Most gRNA-expressing cells are expected to become null or kinase-dead, but at a rare frequency, CDK12 variants that maintain kinase activity can be generated. Subsequent treatment of individual pools with CTX-439 can enrich mutant cells resistant to CTX-439. Out of 29 gRNAs tested, we found surviving colonies from 5 gRNAs. We isolated these colonies, and analyzed their genomic sequence and response to CTX-439 (Supplementary Fig. 11A, B). With an exception of gRNA 18, a single variant was found at each target site, indicating that these are subclones of a single parent. Mutations generated by gRNAs 5, 6 and 7 were closely located, while gRNAs 2 and 18 were located farther apart. IC_50_ values of these resistant colonies were indeed significantly higher than the control gRNA-expressing cells with mutants generated from gRNA 6 (D817N) being the most resistant (IC_50_ ∼1.5 μM). To confirm that these CDK12 variants genuinely caused CTX-439 resistance, we generated gRNA-resistant mutant CDK12 cDNA, expressed them in Cas9-expressing cells, and finally knocked out the endogenous CDK12 gene. These mutant-expressing cells were also significantly resistant (Supplementary Fig. 11C). Strikingly, when 3 distantly located mutations, namely TA770FL, D817N and E1041AG, were combined, cells showed almost complete resistance (Supplementary Fig. 11C). This strongly supports that CTX-439 does not show detrimental off-target inhibition. We next examined the kinase activity of this triple mutant. The p-S2 level in wild-type CDK12-expressing cells showed dose- and time-dependent reductions that were comparable to those in the control cells (Supplementary Fig. 11E-F). In contrast, mutant CDK12-expressing cells maintained the p-S2 levels even in the presence of 1.0 μM CTX-439 (Supplementary Fig. 11E-F). These results indicate that the CDK12 variants identified maintain kinase activity while refractory to CTX-439 treatment.

YM815, D817 and TA770 are located within the ATP pocket which most of the CDK12 inhibitors developed so far use for target binding (Supplementary Fig. 11D), highly suggesting that these variants are gatekeeper mutations.

We then expressed the triple-mutant or wild-type CDK12 in OV90 cells and knocked out endogenous CDK12. The cells showed expected CTD phosphorylation patterns in the absence and presence of CTX-439 (Fig. 6A-B). The mutant cells were highly resistant with no growth suppression up to 1.0 μM (Fig. 6C). We next examined whether CTX-439-mediated MCL1 suppression at both protein and RNA levels was a genuine consequence of CTX-439-mediated inhibition of CDK12/13 function. When empty control and cells expressing wild-type CDK12 were treated with CTX-439, MCL1 and BIM were downregulated and cPARP was induced when AZD4320 was further added (Fig. 6D). In contrast, these CTX-439-associated effects were all canceled in cells expressing mutant CDK12 (Fig. 6D). At the transcript level, *MCL1* upregulation was detected in the nuclear RNA extracted from empty and wild-type CDK12 expressing cells, but no such upregulation was observed in cells expressing mutant CDK12 (Fig. 6E). In addition, readthrough transcription in the *MCL1* and *TUBB4B* genes was detected in empty control and wild-type CDK12 expressing cells, but the degree of readthrough transcription was lower in cells expressing mutant CDK12 (Fig. 6F-G). Taken together, the effects on proliferation as well as protein and RNA expression were caused by specific inhibition of CDK12/13 by CTX-439.

**Fig. 5.**
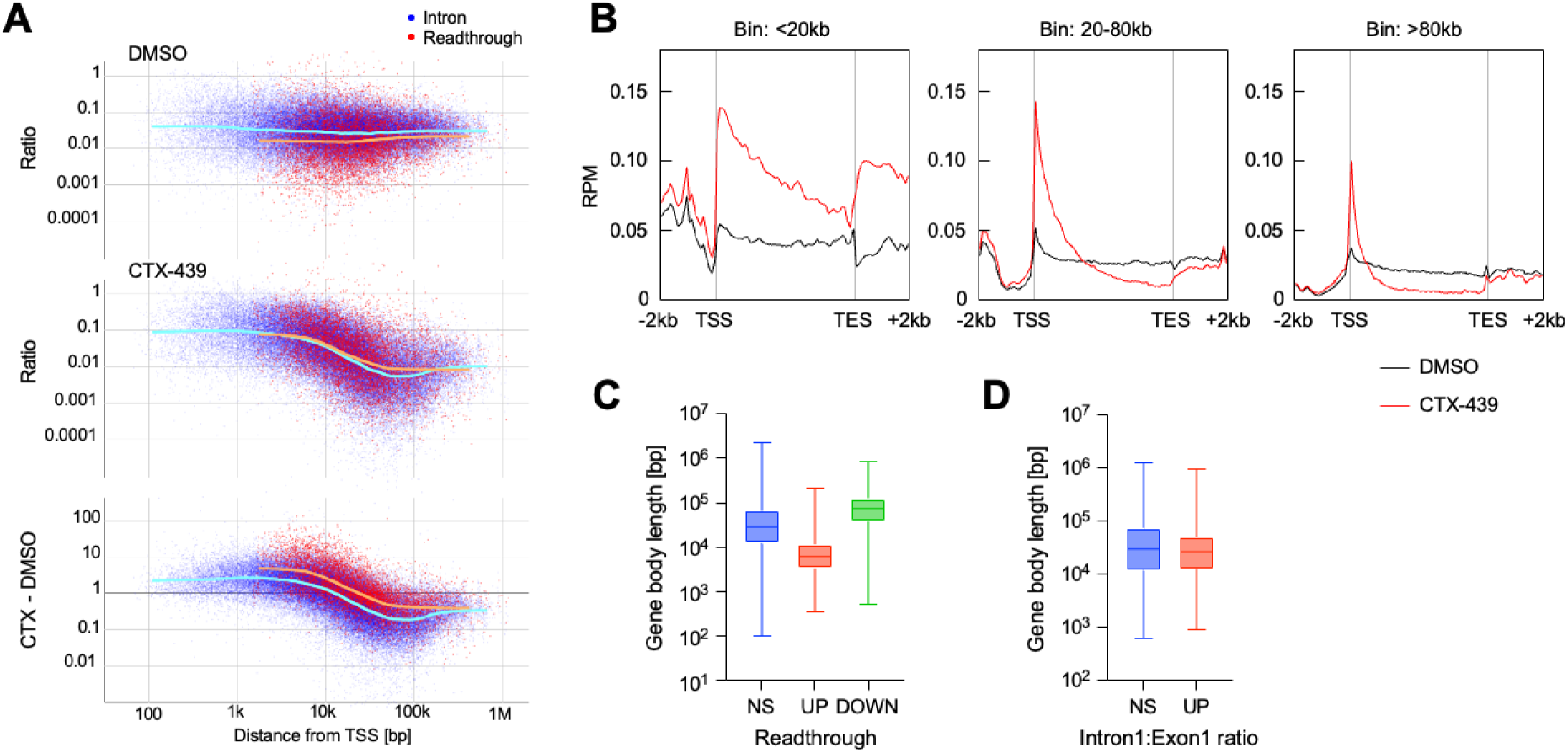
Effects of CTX-439 on transcription is dependent on gene body length. (**A**) Metagene plot showing read density of individual introns (blue) and a 3-kb region immediately after the last exon of each gene (red). The *x* axis represents the distance from TSS to the middle point of each element. RNA-seq data obtained from rRNA-depleted nuclear RNA were analyzed. (**B**) Metagene profile of genes with the indicated gene body length. (**C**) Box plot showing gene body length of genes grouped by differentially expressed readthrough (Figure 3C). (**D**) Box plot showing gene body length of genes grouped by Intron1:Exon1 ratio (Figure 3E). Box plots show median, 25^th^ and 75^th^ percentiles with whiskers of the max and min values.

**Fig. 6.**
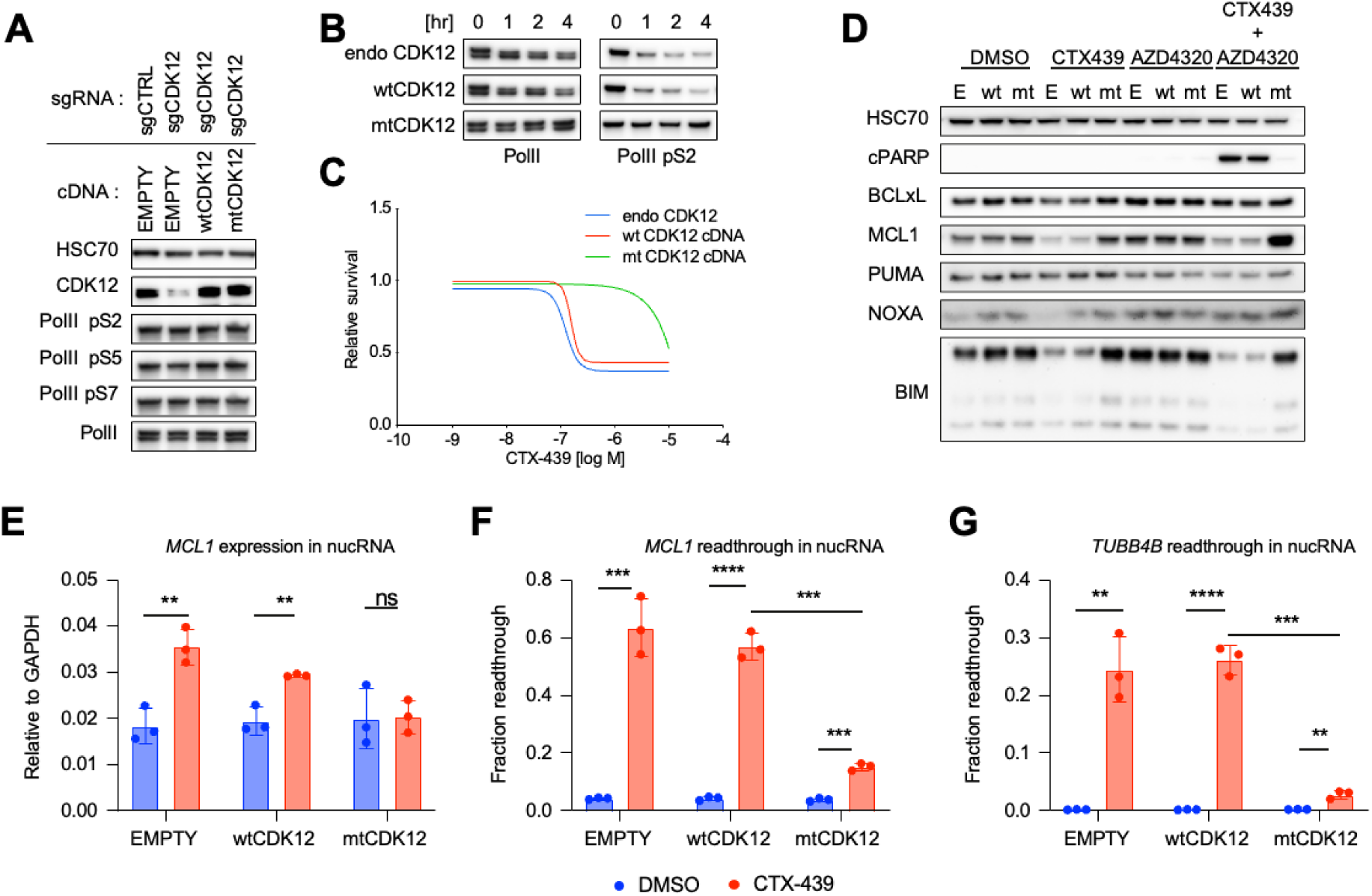
Apoptosis and transcriptional readthrough were directly caused by CTX-439-mediated CDK12/13 inhibition. (**A**) Western blot analysis of cells expressing the indicated cDNA and sgRNA. mtCDK12 carries the combined mutations of TA770FL, D817N and E1041AG. (**B**) Western blot analysis showing that pS2 was maintained in mtCDK12-expressing cells under the treatment of CTX-439. (**C**) Cell survival assay. Cells with endogenous, or exogenous wt or mt CDK12 were tested. (**D**) Western blot analysis. Cells with endogenous (E), or exogenous wt or mt CDK12 were treated with the indicated drug(s) for 8h. Note that MCL1 downregulation was suppressed by mt CDK12. (**E**) RT-qPCR analysis of *MCL1* expression in the nucleus. Note that nuclear *MCL1* transcript upregulation was suppressed by mt CDK12. (**F-G**) RT-qPCR analyses of *MCL1* (**F**) and *TUBB4B* readthrough (**G**) in the nucleus. Readthrough was detected in mt CDK12-expressing cells, but their level is far below than that in cells with endogenous or exogenous wt CDK12. Data are shown as mean ± SD (n=3, **E-G**). * *p*<0.05, ** *p*<0.01, *** *p*<0.001, **** *p*<0.0001 by Student’s *t-*test.

### CDK12/13 and BCL-2/BCL-xL dual inhibition exhibited superior anti-tumor effects in a xenograft model

Lastly, to evaluate the therapeutic potential of dual inhibition of CDK12/13 and BCL-2 family proteins, we examined whether CTX-439 and AZD4320 combination was effective *in vivo*. Firstly, we examined whether *MCL1* readthrough was similarly induced *in vivo* using total RNA extracted from SUM149PT xenografts. As shown in Figure 7A, *MCL1* readthrough transcription was detected as soon as 1h post administration and peaked at 2h, followed by gradual decrease. CDK12 inhibition was observed as shown by the downregulation of self-phosphorylation at S423 and the protein level due to its large gene size (73kb). CTX-439 treatment downregulated the MCL1 protein level, while AZD4320 alone increased the MCL1 level probably due to compensatory effects (Fig. 7B). Consistent with our *in vitro* observation, combination treatment of CTX-439 and AZD4320 were significantly more effective in inducing apoptosis (cPARP and cCasp3), which resulted in greater tumor suppression than either drug alone (Fig. 7C). The body weight did not show significant changes by drug combination during the course of drug treatment (Fig. 7D), suggesting tolerance for treatment. These results support the further development of a combination therapy of CDK12/13 and anti-apoptotic BCL-2 family inhibition.

**Fig. 7.**
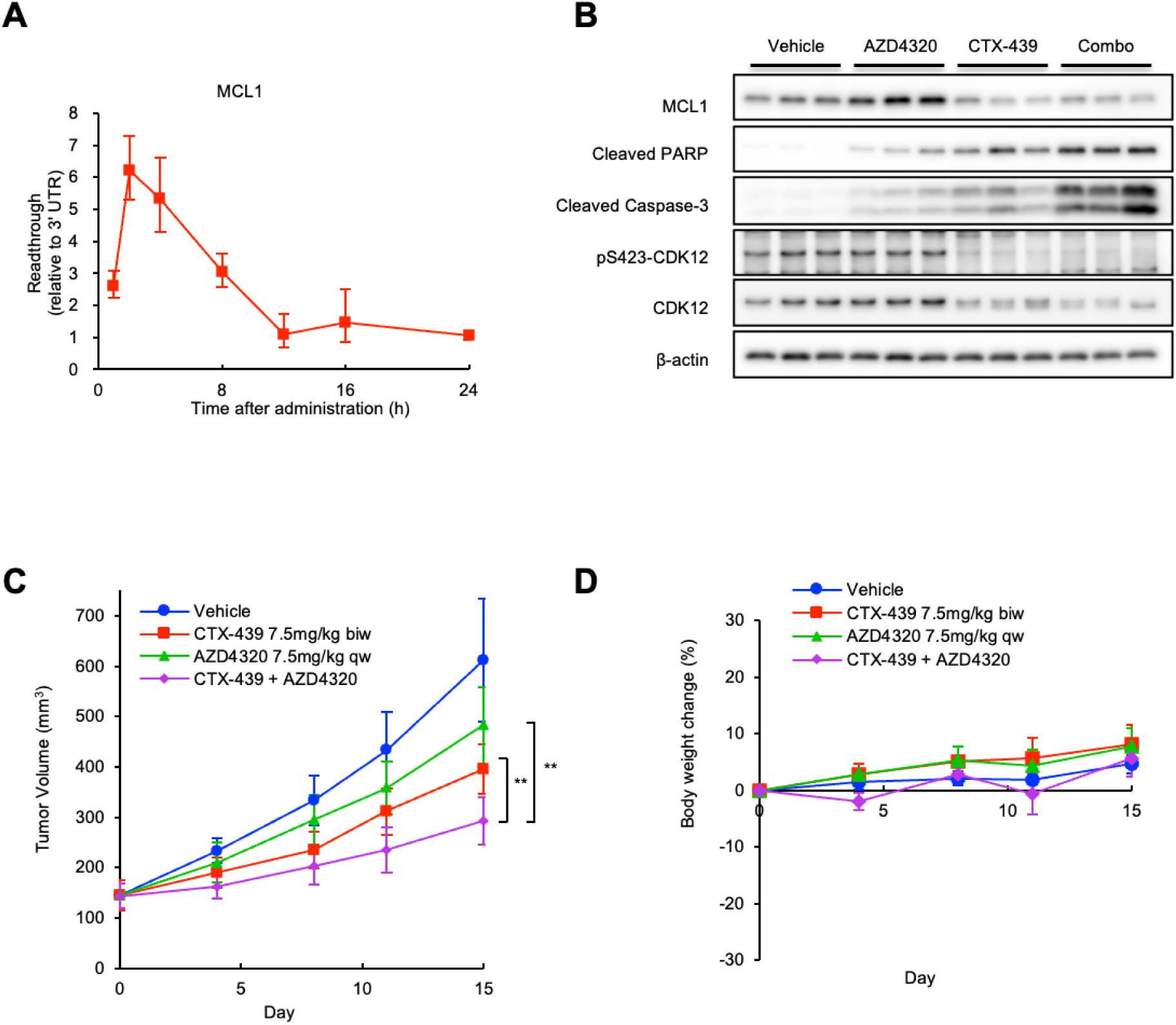
Decreased MCL1 protein expression via *MCL1* read-through and the combination effect of CTX-439 and BCL-2/xL inhibitor AZD4320 in xenograft models. (**A**) RT-qPCR analysis of *MCL1* readthrough in tumors of SUM149PT xenograft models administrated with 30 mg/kg of CTX-439. Tumor samples were collected and RNA was extracted at the indicated time after administration. (**B**) Western blot analysis in tumors of the SUM149PT xenograft model administered with 30 mg/kg of CTX-439 and/or 20 mg/kg of AZD4320. Tumor samples were collected after 8h. (**C**) Enhanced antitumor efficacy of CTX-439 in combination with AZD4320 in the SUM149PT xenograft model. CTX-439 and AZD4320 were administered at the indicated doses and regimens. ** *p*<0.01 by Student’s *t*-test. (**D**) No significant changes in body weight were observed. Data are shown as mean ± SD (n=3, **A** and **B;** n=6, **C** and **D**).

## Discussion

CDK12 and CDK13 have emerged as promising targets for cancer therapy due to their essential roles in transcription regulation and genome stability (Quereda *et al*, 2019; Zhang *et al*, 2016; Iniguez *et al*, 2018). Here, we developed and characterized the novel IND-ready CDK12/13 inhibitor, CTX-439, which possesses favorable physical properties, pharmacokinetics and safety profile in mice, and more importantly, exhibits profound anti-tumor activity. A Phase I clinical trial (NCT06600789) evaluating CT7439, a CDK12/13 inhibitor/Cyclin K glue-degrader, has recently commenced in patients with advanced solid tumors (Ottaviani *et al*, 2025), which underscores a growing interest in this therapeutic strategy. This study demonstrates that CTX-439 offers unique mechanistic insights into transcriptional regulation and highlights its potential in combination with BCL-2 family inhibitors. Our findings enhance the rationale for targeting this pathway by presenting a novel mechanism of action and combination potential.

Recent studies focusing on transcriptional regulation by CDK12 commonly revealed that CDK12 activity is required for transcriptional elongation and that defects cause premature cleavage and polyadenylation, especially for long genes (Dubbury *et al*, 2018; Krajewska *et al*, 2019). Mechanistically, this activity is required to maintain transcription rates and to recruit RNA processing factors such as splicing factors and the U1 snRNAP complex (Krajewska *et al*, 2019), thereby completing transcription until the 3’ end of the gene (Fan *et al*, 2020). This is different from CDK9, which also phosphorylates the same S2 site of Pol II CTD but functions more on the release of Pol II from proximal pausing, thereby promoting productive elongation. CDK9 inhibition therefore leads to the global transcriptional suppression and affects the expression of short-lived transcripts most (Lam *et al*, 2001); sensitivity to CDK9 inhibition is associated with cellular dependency on MYC or MCL1 (Cidado *et al*, 2020; Boffo *et al*, 2018). In contrast, CDK12/13 inhibition affects transcriptional elongation of long genes, and gives a negative impact, particularly, on DDR gene expression (Dubbury *et al*, 2018). CDK12-deficient tumors show characteristic large genomic alterations such as Mb-sized tandem duplication (Li *et al*, 2020), which further indicates that CDK12 is involved in the expression of the DDR genes. As a result of reduced Pol II processivity by CDK12/13 inhibition, a free Pol lI reservoir available for the next round of transcription can increase. This could in turn cause more frequent transcriptional initiation and thus 10-20 kb regions from TSS of every gene can be transcribed more, although transcription beyond this distance is again suppressed due to reduced processivity. This phenomenon was previously captured by TT-seq (Krajewska *et al*, 2019) but our analysis using rRNA-depleted nuclear RNA also revealed a similar result. If gene body size falls within this range, such genes would consequently be upregulated, subject to the efficiencies of splicing and 3’ end processing. Our observations of transcriptional abnormalities induced by CTX-439 thus largely agree with those detected by THZ531 (Krajewska *et al*, 2019).

Nevertheless, our study provides an intriguing insight into the complex regulatory outcome of MCL1 expression by CDK12/13 inhibition. It was previously shown that CDK12/13 inhibition by THZ531 downregulated MCL1 expression at the protein level (Zeng *et al*, 2018), which we also confirmed with CTX-439. However, this observation contradicts the anticipated upregulation for short genes (*MCL1* gene body is 5.1-kb long). In fact, we detected upregulation of *MCL1* expression in the nucleus upon CTX-439 treatment, but this upregulation does not simply translate into cytoplasmic changes. Our results collectively propose the following scenario: the reduction in the efficiency to maturate *MCL1* transcripts by splicing and/or 3’ end processing results in a reduced mRNA supply despite transcriptional upregulation, and then, coupled with its extremely fast mRNA turnover rate, the cytoplasmic *MCL1* mRNA level undergoes downregulation. As MCL1 expression is also regulated post-translationally by the proteasomal degradation, the MCL1 protein level is destined to rapidly decrease. As shown in the RNA-seq analysis using rRNA-depleted nuclear RNAs, which include pre-mRNA molecules (Fig. 3A), *MCL1* transcripts are fairly efficiently processed for both splicing and at the 3’ end as readthrough reads were barely detected in the control cells. However, CDK12/13 inhibition dramatically and acutely disrupts 3’ end processing and transcription termination, leading to long readthrough transcription.

Together with splicing defects, this collectively resulted in the delay in supplying mature mRNA despite transcriptional upregulation. Mature polyA-tailed mRNA with an extended 3’ UTR is likely to be produced, but such mRNA may not be transported efficiently to the cytoplasm as shown by the strong discrepancy of *TUBB4B* readthrough between the nucleus and the cytoplasm (Supplementary Fig. 8G). In addition, we recently identified mRNA transport factor NXF1 as a CDK12-interacting protein (Ebara *et al*, 2024) and thus this pathway may also be directly affected by CDK12/13 inhibition. This scenario is also consistent with the observation that the replacement of the native *MCL1* 3’ UTR with a more efficient bpA sequence restores the consistency of transcriptomic changes in the nucleus and the cytoplasm, leading to MCL1 upregulation at the protein level. This indicates that the splicing defect seen in the *MCL1* intron 1 contributes less to downregulation. Taken together, MCL1 responds uniquely to CDK12/13 inhibition and the mechanism of its downregulation overtly differs from that of CDK9 inhibition.

MCL1 is associated with a poor prognosis and is a potential therapeutic target in breast cancer, particularly in highly aggressive TNBC (Campbell *et al*, 2018). Moreover, MCL1 has been recognized as a key survival factor in various solid and hematologic malignancies, underscoring its biological significance in MCL1-driven tumors (Kotschy *et al*, 2016; Caenepeel *et al*, 2018). These findings support the rationale for targeting MCL1 and highlight the potential of CTX-439 against TNBC and other MCL1-driven cancers. Future research is needed to elucidate the molecular determinants of response, including MCL1 expression and associated apoptotic pathways, to refine patient selection strategies and maximize clinical benefit.

Gatekeeper mutations are a common issue that cause drug resistance against kinase inhibitors. For instance, a threonine to isoleucine substitution at position 315 in the ABL part of BCR-ABL is sufficient to become resistant to imatinib, which led to the development of 2nd-generation ABL inhibitor, ponatinib (O’Hare *et al*, 2009). Another example is the T790M mutation in EGFR, which renders resistance to gefitinib and erlotinib (Yun *et al*, 2008) and led to the development of osimertinib, active against T790M mutant(Cross *et al*, 2014). CTX-439 is an ATP-competitive inhibitor and thus may be inactivated by similar resistance mechanisms. With a limited number of gRNAs screened, we successfully identified 5 variants within the kinase domain that show decreased response.

Mapping these variants on the structure of the CDK12 kinase domain revealed that they are indeed located in the ATP binding groove, where CTX-439 most likely binds. Structural analysis of the CTX-439-bound form of the CDK12 kinase domain could clarify the basis of this resistance. While not all mutations may be clinically relevant, cataloging potential resistance variants can facilitate rapid assessment in future clinical settings. Notably, cells expressing mutant CDK12 showed diminished response to CTX-439. This suggests that the observed anti-proliferation effect is directly induced by CTX-439 and that off-target effects are negligible. These findings definitively confirm the compound’s on-target specificity and support its further development as a therapeutic agent.

In summary, we developed CTX-439, a novel and highly specific CDK12/13 inhibitor that exhibits robust anti-tumor activity in preclinical models. While clinical data are not yet available, our results provide a strong rationale for advancing CTX-439 toward clinical evaluation. While CTX-439 demonstrates efficacy as monotherapy, our data suggest that BCL-2/xL expression can modulate therapeutic response. In this context, combination strategies involving dual inhibition of CDK12/13 and anti-apoptotic factors could enhance treatment outcomes and present a promising direction for future clinical investigation. These insights highlight the potential of CDK12/13 inhibition as a versatile and impactful approach to cancer therapy.

## Methods

### Compounds

CTX-439 is prepared as described below. THZ531, SR-4835, AZD4320, AZD5991, and A-1331852 were purchased from MedChem Express or Selleck.

### Synthesis of CTX-439 General chemical procedure

^1^H NMR spectra were recorded on Bruker AVANCE III (300 MHz), Chemical shifts are given in parts per million (ppm) downfield from tetramethysilane (δ) as the internal standard in deuterated solvent and coupling constants (J) are in Hertz (Hz). Data are reported as follows: chemical shift, integration, multiplicity (s = singlet, d = doublet, t = triplet, q = quartet, m = multiplet, dd = doublet of doublet, brs = broad singlet) and coupling constants. All solvents and reagents were obtained from commercial suppliers and used without further purification. LC-MS analysis was performed on Shimadzu UFLC-Mass Spectrometer System, operating in or ESI (+ or −) ionization mode. Analyses were eluted using a linear gradient of 0.05% TFA containing water/acetonitrile, 0.1% TFA containing water/acetonitrile or 5 mM ammonium acetate containing water/acetonitrile mobile phase.

### Synthetic procedures

#### 4-(5-(methylsulfonyl)pyridin-3-yl)-2-(methylthio)-5-(trifluoromethyl)pyrimidine (3)

A mixture of 4-chloro-2-(methylthio)-5-(trifluoromethyl)pyrimidine (**1**, 1.0 g, 26.2 mmol), 3-(methylsulfonyl)-5-(4,4,5,5-tetramethyl-1,3,2-dioxaborolan-2-yl)pyridine (**2**, 1.3 g, 27.5 mmol), PdCl_2_(Amphos)_2_ (310 mg, 0.44 mmol), 2 M sodium carbonate aqueous solution (4.4 mL) and DME (10 mL) was stirred under microwave irradiation at 100 °C for 1 h. A similar reaction was repeated 6 times (in other words, 6 batches were obtained). To the combined reaction mixture was added water at room temperature, and the mixture was extracted with ethyl acetate. The organic layer was separated, washed with saturated brine, and dried over anhydrous magnesium sulfate. The solvent was evaporated under reduced pressure. The residue was purified by silica gel column chromatography (ethyl acetate/hexane) to give the title compound (7.1 g, 77%). MS: [M+H]^+^ 350.0.

#### 2-(methylsulfonyl)-4-(5-(methylsulfonyl)pyridin-3-yl)-5-(trifluoromethyl)pyrimidine (4)

To a solution of **3** (7.1 g, 20.3 mmol) in ethyl acetate (200 mL) *m*-chloroperbenzoic acid (11.0 g, ca 70%, 44.6 mmol) was added at 0 °C. After stirring at room temperature for 5 h, the reaction solvent was evaporated under reduced pressure. The residue was purified by silica gel column chromatography (ethyl acetate/hexane) to give the title compound (5.7 g, 74%). MS: [M+H]^+^ 382.0.

#### *tert*-butyl (*trans*-4-((5-bromopyrazin-2-yl)amino)cyclohexyl)carbamate (7)

To a mixture of *tert*-butyl (*trans*-4-aminocyclohexyl)carbamate (**6**, 20 g, 93.3 mmol) and 1-butanol (100 mL) 2,5-dibromopyrazine (**5**, 26.6 g, 112 mmol) and DIPEA (24.1 g, 142 mmol) were added, and the mixture was stirred under reflux overnight. To the reaction mixture water was added, and the mixture was extracted with ethyl acetate. The organic layer was dried over anhydrous sodium sulfate, and the solvent was evaporated under reduced pressure. The residue was purified by silica gel column chromatography (ethyl acetate/petroleum ether) to give the title compound (22.8 g, 66%). MS: [M+H-tBu]^+^ 315.1.

#### *tert*-butyl (*trans*-4-((5-(2-methoxypyrimidin-5-yl)pyrazin-2-yl)amino)cyclohexyl)carbamate (9)

A mixture of **7** (1.5 g, 4.04 mmol), (2-methoxypyrimidin-5-yl)boronic acid (**8**, 0.933 g, 6.06 mmol), PdCl_2_(dppf) (0.302 g, 0.37 mmol), 2 M cesium carbonate aqueous solution (5.05 mL) and DME (20 mL) was stirred under a nitrogen atmosphere at 90 °C for 2 h. Water was added to the reaction mixture at room temperature, and the mixture was extracted with ethyl acetate and THF. The organic layer was separated, washed with saturated brine, and dried over anhydrous magnesium sulfate. The solvent was evaporated under reduced pressure. The residue was purified by silica gel column chromatography (NH, ethyl acetate/hexane) to give the title compound (1.41 g, 87%). MS: [M+H]^+^ 401.2.

#### *tert*-butyl (*trans*-4-(1-(5-(2-methoxypyrimidin-5-yl)pyrazin-2-yl)butylamido)cyclohexyl)carbamate (11)

To a mixture of **9** (1 g, 2.50 mmol), DMAP (275 mg, 2.25 mmol) and DMA (30 mL) butyryl chloride (**10**, 1.3 mL, 12.4 mmol) was slowly added at room temperature. The mixture was stirred at 70 °C for 4 h, and at 100 °C for 2 h. The reaction mixture was added to water at room temperature, and the mixture was extracted with ethyl acetate. The organic layer was separated, washed with water and saturated brine, and dried over anhydrous sodium sulfate. The solvent was evaporated under reduced pressure. The residue was purified by silica gel column chromatography (ethyl acetate/hexane) to give the title compound (214 mg, 18%).

MS: [M+H]^+^ 471.3.

#### *N*-(*trans*-4-aminocyclohexyl)-*N*-(5-(2-methoxypyrimidin-5-yl)pyrazin-2-yl)butylamide (12)

A mixture of **11** (214 mg, 0.45 mmol) and TFA (2 mL) was stirred at room temperature for 1 h. The reaction mixture was concentrated under reduced pressure, and the residue was purified by silica gel column chromatography (NH, methanol/ethyl acetate) to give the title compound (144 mg, 85%).

#### *N*-(*trans*-4-((4-(5-(methanesulfonyl)pyridin-3-yl)-5-(trifluoromethyl)pyrimidin-2-yl)amino)cyclohexyl)-*N*-(5-(2-methoxypyrimidin-5-yl)pyrazin-2-yl)butanamide (CTX-439)

A mixture of **12** (144 mg, 0.39 mmol), **4** (163 mg, 0.43 mmol), DIPEA (0.2 mL, 1.18 mmol) and DMF (2 mL) was stirred at 70 °C for 3 h. The reaction mixture was added to water at room temperature, and the mixture was extracted with ethyl acetate. The organic layer was separated, washed with water and saturated brine, and dried over anhydrous sodium sulfate. The solvent was evaporated under reduced pressure. The residue was purified by silica gel column chromatography (ethyl acetate/hexane and NH, ethyl acetate/hexane) to give the title compound (66 mg, 25%). ^1^H NMR (300 MHz, DMSO-*d*_6_) δ 0.73-0.82 (3H, m), 1.20-1.57 (6H, m), 1.79-2.02 (6H, m), 3.32-3.42 (3H, m), 3.66 (1H, br s), 3.98-4.07 (3H, m), 4.43 (1H, br s), 8.19-8.48 (2H, m), 8.68-8.78 (2H, m), 8.96-9.02 (1H, m), 9.15-9.22 (1H, m), 9.25-9.39 (3H, m). MS: [M+H]^+^ 672.3.

### Kinome Profiling

Kinase selectivity was assessed based on affinity and inhibitory activity against kinases. Affinity to CDKs was determined using DiscoverX KinomeScan to determine Kd values, and affinity to 468 kinases was screened using the ScanMAX kinase panel at a concentration of 30 nM, which is 100 times the Kd value of CDK12. Kinases with high affinity (percentage control ≤ 30) were re-assayed and Kd values were determined.

Compound was tested in duplicate at 11 doses of serial 3-fold dilutions starting at 10 μM. The detailed protocol is available from DiscoverX. Kinome trees were constructed based on these Kd values (Eid *et al*, 2017). The figure is reprinted with permission from Cell Signaling Technology, Inc. Inhibitory activity against kinase activity was evaluated using the HotSpot assay from Reaction Biology. Compounds were tested in duplicate at 10 doses of serial 3-fold dilutions starting at 10 μM. Reactions were carried out at Km ATP. Kinase activity data were expressed as the percent remaining kinase activity in test samples compared to vehicle (DMSO) reactions. IC_50_ values and curve fits were obtained using Prism4 Software (GraphPad). Ki values were calculated from IC_50_ values using the Cheng and Prusoff equation.

### Cell culture

Breast cancer cell line, SUM149PT, was purchased from BioIVT and cultured in Ham’s F12 (Thermo Fisher) supplemented with 5% fetal bovine serum (FBS), 10 mM HEPES, 1 μg/mL hydrocortisone and 5 μg/mL insulin as instructed. OV90 was purchased from ATCC and cultured in DMEM/F12 (Thermo Fisher) supplemented with 10% FBS. Breast cancer cell lines (Supplementary Table 2), ovarian cancer cell lines (Supplementary Table 2), pancreatic cancer cell lines (Capan1 and MiaPaca2) and colorectal cancer cell lines (HCT116, SW620 and RKO) were described before (Behan *et al*, 2019). 293FT was cultured in DMEM (Wako, 041-30081) supplemented with 10% FBS. All cell lines tested negative for mycoplasma.

### Plasmids

For CRISPR-a, pKLV2-EF1adCas9VP64T2ABsd-W (Addgene #112919) and pKLV2-EF1aMS2p65HSF1hyg-W (Addgene #112920) were used. For CRISPR-KO, pKLV2-EF1a-Cas9Bsd-W (Addgene #68343), pKLV2-U6gRNA5(BbsI)-PGKpuro2ABFP-W (Addgene #67974) and pKLV2.2-h7SKgRNA5(SapI)-hU6gRNA5(BbsI)-PGKpuroBFP-W (Addgene #72666) were used.

To construct a HA-tagged MCL1 expression vector, *MCL1* cDNA was first generated by RT-PCR and cloned into the AscI site of pKLV2-EF1a-IRESneo (Addgene #234503), resulting in pKLV2-EF1a-MCL1HA-IRESneo (Addgene #234504).

To construct HA-tagged CDK12 expression vectors, *CDK12* cDNA fragment was first generated by PCR using pHAGE-CDK12 (Addgene #116723) as a template. During this process, silent mutations, which remove CRISPR sites (gCDK12_v3_6-2 and gCDK12_v3_6-4), were added. The fragment was then cloned into pKLV2-EFa-IRESneo, resulting in pKLV2-EF1a-HACDK12wt-IRESneo (Addgene #234505). Subsequently, the mutations identified in g2, g5, g6, g7 and g18 were introduced into the gRNA-resistant CDK12 coding region by site-directed mutagenesis (Addgene #234506-10). A single cDNA carrying three mutations (g2, g6 and g18) was also generated (Addgene #234511). PCR was performed using Q5 Hot Start HiFi PCR master mix (NEB). All cloning was performed using the NEBuilder HiFi DNA Assembly master mix (NEB). PCR-amplified regions were verified by Sanger sequencing. Primers were listed in Supplementary Table 6.

### CRISPR-activation library

The gRNA expression cassette of the Calabrese library (Addgene #92379 and #92380) was PCR-amplified and cloned into the MluI/BamHI site of pKLV3-U6gRNA5(BbsI)-PGKpuroBFP-W-L1 using NEBuilder Assembly. The assembly mixture was purified using the MinElute PCR purification kit (Qiagen) and electroporated into NEB10-beta (NEB). Sets A and B were separately cloned and verified by deep sequencing. These sets were combined prior to virus production. Primers were listed in Supplementary Table 6.

### Lentivirus production and transduction

293FT cells were pre-seeded at 9.6 x 10^5^ cells per well of 6-well plates. On the following day, transfer vector (0.9 μg), psPax2 (0.9 μg), pMD2.G (0.2 μg) and PLUS reagent (2 μL) were added to 500 μL of Opti-MEM and incubated for 5 min. LipofectamineLTX (6 μL) was added and incubated for 30 min. DNA mixture was then added to 293FT. After 16 h incubation, medium was replaced with fresh medium and cells were further incubated for 1 day. Viral supernatant was filtered using a 0.4 μm syringe filter and then stored at -80 °C. Transduction was carried out by simultaneously adding target cells and viral supernatant in the presence of 8 μg per mL polybrene into appropriate culture plates. Virus was removed on the following day by refresh culture media. Two-to-three days post transduction, cells were trypsinized and analyzed by cytometry for BFP expression in case of gRNA expression. For knockout purposes, 20-60% BFP-positivity was accepted and the cells were subjected to puromycin selection.

### CRISPR-activation screen

To generate a CRISPR-a cell line, SUM149PT was transduced with dCas9-VP64 and selected with Blasticidin, followed by transduction with SM2p65HSF1 activation effectors and hygromycin selection. 3 x 10^7^ CRISPR-a cells were transduced with the CRISPR-a library at a transduction rate of 30 % (by BFP positive) and selected with puromycin. On day 13, 4-5 x 10^7^ cells were seeded onto three T525 flasks. On the following day, CTX-439 was added at a concentration of 150 nM and the cells were treated for 8 days. Drug-containing medium was refreshed on day 4. In parallel, DMSO-control culture was similarly prepared. On day 13, ∼3 x 10^7^ cells were harvested as a day 0 control. Genomic DNA extraction and gRNA amplification were performed as described previously (Behan *et al*, 2019; Tzelepis *et al*, 2016). Sequencing libraries were analyzed by DNBSEQ-G400RS with a custom sequencing primer (Behan *et al*, 2019; Tzelepis *et al*, 2016).

### CRISPR screening data analysis

Statistical analysis was performed using MAGeCK package (Li *et al*, 2014). Two comparisons were performed: 1. Day 0 vs DMSO day 8, and 2. Day 0 vs CTX-439 day 8. Resistant hits were identified with the following filterings: LFC > 0.5 in Day 0 vs CTX-439, LFC < 0.5 in Day 0 vs DMSO, and positive p-value < 0.01 in Day 0 vs CTX-439.

### Isolation of CDK12 variants with CTX-439 resistance

Twenty nine gRNAs targeting kinase domain coding exons of CDK12 (Supplementary Table 5) were designed and cloned individually into pKLV2-U6gRNA5(BbsI)-PGKpuroBFP-W. Lentiviruses were produced as described above. Cas9-expressing MDA-MB-436 (3 x 10^5^ cells) were transduced with the virus individually in a transduction rate of ∼30%, which yielded ∼1 x 10^5^ non-redundant mutants, and selected in 3 μg/mL puromycin. 3 x 10^6^ selected cells were seeded on a 15-cm dish and on the following day CTX-439 was added at 200 nM. Approximately 2 weeks later, resistant colonies were picked (4 colonies each) and expanded. Genomic DNA was extracted using DNeasy Blood and Tissue kit (Qiagen) and a region surrounding the gRNA target site was PCR-amplified. Because direct Sanger sequencing resulted in mixed sequence signals, PCR products were cloned into a plasmid and then 10 clones each were subjected to Sanger sequencing.

### Drug response assay

A thousand cells were seeded in each well of 96-well plates. On the following day, a test compound was serially diluted (typically, 2-fold dilution starting from 10 μM) and added to each well with uniform DMSO concentration across conditions. Cells were cultured for 3 days and viability was assessed using CellTiter-Glo (Promega) according to the manufacturer’s protocol.

### Crystal violet staining

1-2 x 10^6^ cells were plated on one 6-well plate. On the following day, cells were treated with drugs at predetermined concentration for 8 days. The plates were washed with PBS once and cells were stained with crystal violet (Nakarai) for 10-15 min. The cells were washed with water and air dried.

### Western blot analysis

Cell lysates were extracted using RIPA buffer (Sigma) with Halt Protease & Phosphatase inhibitor cocktail (Thermo Fisher) and protein concentration was determined using the Pierce 660nM Protein Assay (Thermo Fisher). 1-10 μg of protein were separated on NuPAGE 4-12 % Bis-Tris gel (Thermo Fisher) and transferred to an Immobilon-P membrane (Millipore). The membrane was blocked with TBS-T (Santa Cruz) with 3% skim milk (Nacalai) or 5% BSA (Wako) for 1 h and treated with a primary antibody overnight at 4 °C. On the following day, the membrane was washed with TBS-T and treated with secondary antibody for 1 h. After washing, signals were detected using SuperSignal West Pico PLUS Chemiluminescent Substrate (Thermo Fisher) with ImageQuant800 (Cytiva). Antibodies used are listed in Supplementary Table 8.

### RNA extraction

Total RNA was extracted using the RNeasy Mini kit (Qiagen) according to the manufacturer’s protocol. Nuclear and cytoplasmic RNA fractionation was carried out using the RNA Subcellular Isolation Kit (Active Motif). For tumor tissues, tumor fragments were homogenized in chloroform, and the supernatants were used to extract RNA using the RNeasy Mini Kit (Qiagen).

### RT-qPCR

Total, nuclear or cytoplasmic RNA (250 ng) was reverse transcribed in a 10-uL reaction volume using SuperScript IV VILO Master Mix (Thermo Fisher). The mixture was then diluted 20-fold with water. Two microliters of the diluted mixture were used for quantitative PCR using PowerUp SYBR Green Master Mix (Thermo Fisher) with QuantStudio (Thermo Fisher). Quantification was performed in duplicate for each primer pair. The ΔΔCt method was used for normalization. Primers are listed on Supplementary Table 7.

### RNA-seq

Total or subcellular RNA was further purified with NEBNext Poly(A) mRNA Magnetic Isolation Module (NEB). Nuclear RNA extracted from OV90 was also purified with the NEBNext rRNA Depletion Kit v2 (NEB) to remove ribosomal RNA. RNA-seq libraries were then generated using the NEBNext Ultra II Directional RNA Library Prep Kit for Illumina (NEB) according to the manufacturer’s instructions. Resulting libraries were sequenced using DNBSEQ-G400RS by 100-bp paired-end sequencing.

### RNA-seq data analysis

Sequencing reads derived from SUM149PT (polyA+) were mapped to the human reference genome hg38 using HISAT2 (v2.2.1). The reads assigned to genes were counted using featureCounts (v2.0.3). Differentially expressed genes were analyzed using DESeq2 (v1.38.2) while filtering out genes with low expression (average read counts in all conditions are less than 10). GO term (Biological process) enrichment analysis was performed with top 3000 differentially expressed genes (LFC < -1, adjusted p-value < 0.01) in ShinyGO 0.82.

Sequencing reads derived from OV90 (polyA+ and rRNA-depleted RNA) were mapped to the human reference genome hg38 using HISAT2 (v2.2.1) and two separate bam files (according to strands) were generated.

3’ readthrough was annotated as follows: First, we defined a readthrough region as a 1-kb region immediately after the last exon of each of the MANE select genes. When two genes are adjacently located in the same transcriptional orientation, a readthrough region that overlaps with exons of the downstream gene was discarded from the analysis (see Supplementary Fig. 8A). Strand-specific gtf files including exons, introns and readthrough regions were separately generated and used to count reads from the bam file with the matched strand using TMACalculator (v0.0.3). By this way, 2 adjacent genes with an inverted orientation can be distinguished (exemplified in Supplementary Fig. 8B).

Differentially expressed readthrough was then analyzed using DEseq2 (v1.38.2). For Figure 3E and Supplementary Figure 9, the mean of per-base read depth for each intron 1 and exon 1 was first calculated, and then the ratio of the intron to the exon was calculated for each gene. Statistical analysis was performed by Welch’s t test.

For Figure 5, the ratio of the mean per-base read depth of each intron or readthrough to that of the whole genome was calculated. In this analysis, readthrough was defined as a 3-kb region immediately after the last exon. Distance from TSS was calculated as a distance between the TSS and the midpoint of each intron or readthrough.

### Animal experiments

Animal studies were approved by the Animal Research Committee of Kyoto University (Y-22-1 and Med Kyo 21538, 22287, 23251). Procedures for the care and use of animals at Chordia Therapeutics Inc. were reviewed and approved by the Institutional Animal Care and Use Committee (IACUC) of the Shonan Health Innovation Park. The care and use of animals was conducted in accordance with the regulations of the Association for Assessment and Accreditation of Laboratory Animal Care International (AAALAC).

### SUM149PT xenograft model

A total of 3 x 10^6^ SUM149PT cells in 100 μl of a 1:1 mixture of HBSS (Thermo) and Matrigel Matrix (BD Biosciences, #14175095) were subcutaneously injected into the flank of 6-week old BALB/cA Jcl-nu/nu mice (CLEA Japan). Tumor growth was monitored with vernier calipers and tumor volume was calculated using the formula (0.5 x [length x width^2^]). When tumor volume reached approximately 200 mm^3^, mice were randomly assigned into treatment groups (n=6/group) and treated with vehicle or CTX-439 (7.5, 15 and 30 mg/kg) via oral gavage (p.o.) twice weekly for 15 days. In the drug combination study, mice were treated with vehicle, CTX-439 (7.5 mg/kg, p.o., twice weekly), AZD4320 (7.5 mg/kg, intraperitoneally (i.p.), once weekly), or in combination. The dosing volume for each mouse was 10 mL/kg. Tumor volume and body weight were measured twice per week. The vehicle for CTX-439 was a mixture of DMSO (WAKO), Cremophor EL (Sigma), propylene glycol (WAKO), and water (Otsuka) (10:10:10:70 [v/v/v/v]). A 10X concentrated DMSO solution of CTX-439 was prepared beforehand; immediately prior to administration, the DMSO solution was thoroughly mixed with an equal amount of Cremophor EL, propylene glycol, and 7X water in sequence. The vehicle for AZD4320 was a mixture of DMSO, PEG300 (MCE), Tween-80 (MCE), and saline (Otsuka) (10:40:5:45 [v/v/v/v]). A 10X concentrated DMSO solution of AZD4320 was prepared, and the DMSO mixture was added to PEG300, Tween-80, and saline and mixed in this order immediately prior to administration.

### Patient-derived xenograft models

Patient characterization of breast cancer PDX models used in this study are described in Supplementary Tables 3 and 4. To generate PDX models, frozen samples were cut into small pieces and transplanted into NOG mice (NOD.Cg-*Prkdc^scid^Il2rg^tm1Sug^*/ShiJic). When tumor volume reached approximately 50-100 mm^3^, mice were randomly assigned and treated with vehicle or CTX-439 (15 mg/kg) via oral gavage twice weekly. All patients included in this study provided written informed consent under the Kyoto University Ethical Approval G0424-20. In addition, two TNBC PDX models were tested by Crown Biosciences, Inc.

### Pharmacokinetic and Pharmacodynamics Experiments

A total of 3 x 10^6^ SUM149PT cells in 100 μL of a 1:1 mixture of HBSS (Thermo) and Matrigel Matrix (BD Biosciences, #14175095) were subcutaneously injected into the flank of 6-week old BALB/cA Jcl-nu/nu mice (CLEA Japan). Tumor growth was monitored with vernier calipers and tumor volume was calculated using the formula (0.5 x [length x width^2^]). When tumor volume reached approximately 200 mm^3^, mice were randomly assigned into treatment groups (n = 3/group) and administered once with vehicle or CTX-439 (7.5, 15 and 30 mg/kg) via oral gavage under ad libitum feeding conditions. Blood and tumor samples were collected at 1, 2, 4, 8, 12, 16 and 24 h after administration under isoflurane anesthesia. Whole blood samples were stored in tubes containing heparin sodium and placed on ice until being centrifuged at the speed of 8,000 rpm for 1 minute. The resulting plasma was separated and stored at -80°C until pharmacokinetic analysis. Tumor fragments were frozen on dry ice and stored at -80°C until pharmacokinetic and pharmacodynamic analysis was performed.

### PK analysis

CTX-439 was dissolved and diluted in DMSO/acetonitrile (2:8) to prepare standard working solutions at 0.0003 to 100 μg/mL. Calibration standard plasma samples were prepared by mixing blank plasma with standard working solution. Plasma samples were mixed with acetonitrile containing internal standard diclofenac, centrifuged, and the supernatants were diluted for liquid chromatography with tandem mass spectrometry (LC/MS/MS) analysis. Calibration standard tumor homogenate samples were prepared by mixing blank tumor homogenate with standard working solution. Tumor samples were homogenized with saline to approximately 20% (w/w) and mixed with acetonitrile containing diclofenac, centrifuged, and the supernatants were diluted for LC/MS/MS analysis.

Calibration curves were evaluated within the concentration ranges of 0.0003 to 10 μg/mL for plasma and 0.005 to 15 μg/g for tumor homogenate. Concentrations of CTX-439 in plasma and tumor were determined by LC/MS/MS at Axcelead Drug Discovery Partners, Inc.

### PD analysis

Pharmacodynamics were analyzed by quantifying the expression of *BRCA1/2* as an established downstream target gene of CDK12/13 inhibition. Total RNA were extracted from the stored tumor fragments and used for RT-qPCR analysis as described above.

### Statistical Analysis

Statistical analyses were performed using GraphPad Prism or Excel. Data are shown as mean ± SD. Significance was determined using the appropriate test as specified in each Figure legend. Statistical significance is denoted as *p<0.05, **p<0.01, ***p<0.001.

## Data availability

All data needed to evaluate the conclusions in the paper are present in the paper and/or the Supplementary Materials. RNA-seq data are available from Gene Expression Omnibus (GSE291144). Please access the data with token ydclmggetravzon

## Author contributions

Conceptualization: DM, KY. Investigation: HY, RK, AM, SE, TH, ST, YT, MS, SS, MT, HT, TT, HB, YA, YF, YF, SI. Supervision: MT, KK, DM, KY. Writing-original draft: HY, KY. Writing-review and editing: HY, YT, KK, DM, KY.

## Acknowledgements

We thank K. Yamamoto for valuable advice and analysis on RNA-seq data. We are also grateful to K. Shimokawa for providing the description of the synthesis method. We gratefully acknowledge A. Shibahara for her support in data organization and mouse husbandry.

This work was supported by the Japan Agency for Medical Research and Development (AMED) under grant number 21ck0106648h (MT, KY and DM).

