## Supplementary Materials for "CDK12/13 inhibitor, CTX-439, suppresses tumor growth and potentiates BCL-2 family blockade"

Hiroko Yamakawa *et al.*

**This PDF file includes:**

Figs. S1 to S11
Tables S1 to S8
Data S1 to S3

#### Figure S1

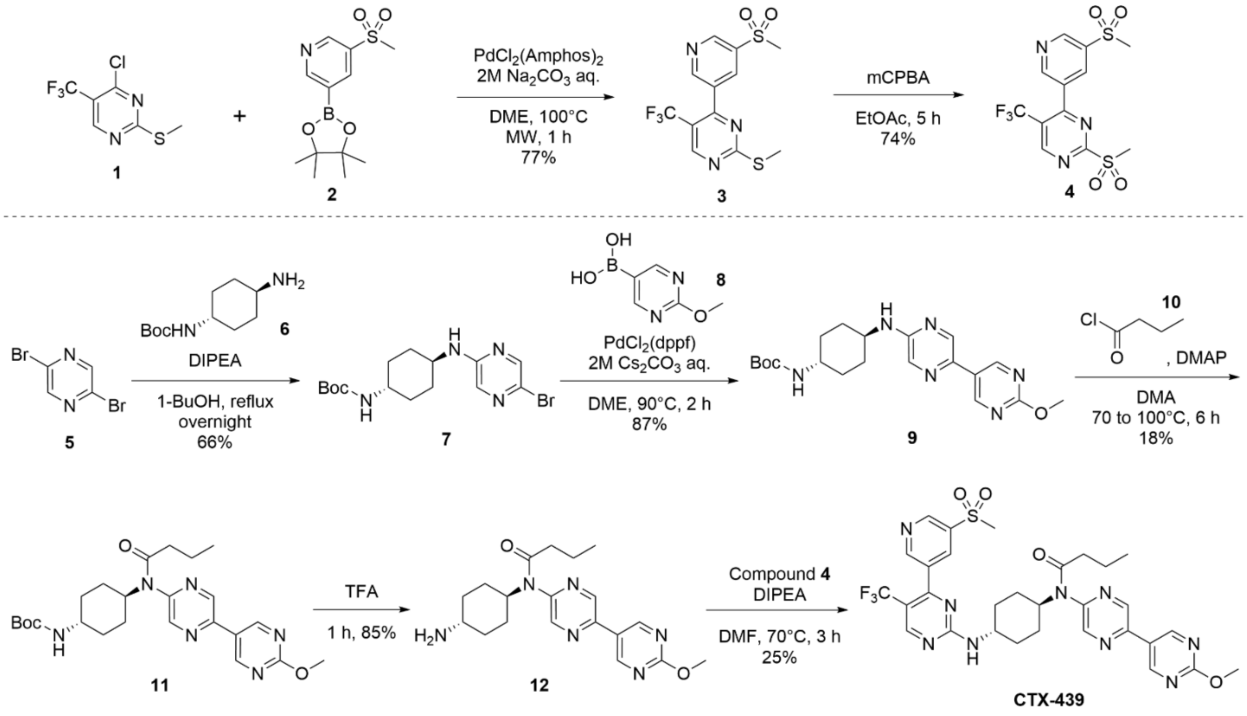


**Figure S1. Synthesis of CTX-439.**
Detailed chemical synthesis procedure was described in Methods. Abbreviations: Boc, tert-butoxycarbonyl; 1-BuOH, 1-butanol; mCPBA, *m*-chloroperbenzoic acid; DIPEA, *N,N*-diisopropylethylamine; DMA, *N,N*-dimethylacetamide; DMAP, 4-dimethylaminopyridine; DME, 1,2-dimethoxyethane; DMF, *N,N*-dimethylformamide; EtOAc, ethyl acetate; $PdCl_2(Amphos)_2$, bis(di-*tert*-butyl(4-dimethylaminophenyl)phosphine)dichloropalladium(II); $PdCl_2$(dppf), [1,1’-bis(diphenylphosphino)ferrocene]dichloropalladium(II); TFA, trifluoroacetic acid.

### Figure S2

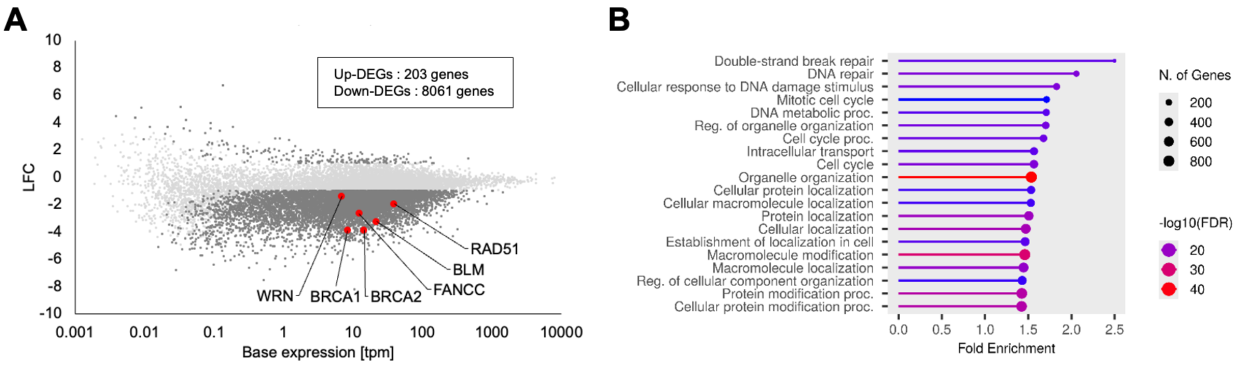


**Figure S2. RNA-seq analysis of CTX-439 treated SUM149PT.**
**A**. MA plot showing global effects on transcription by CTX-439. Cells were treated with 100 nM CTX-439 for 6h in triplicate. Significantly altered genes (*padj*<0.01, |LFC|>1) were highlighted in dark grey. DDR genes were also highlighted by red. **B**. GO term (Biological process) enrichment of downregulated genes (top3000 ranked by LFC).

#### Figure S3

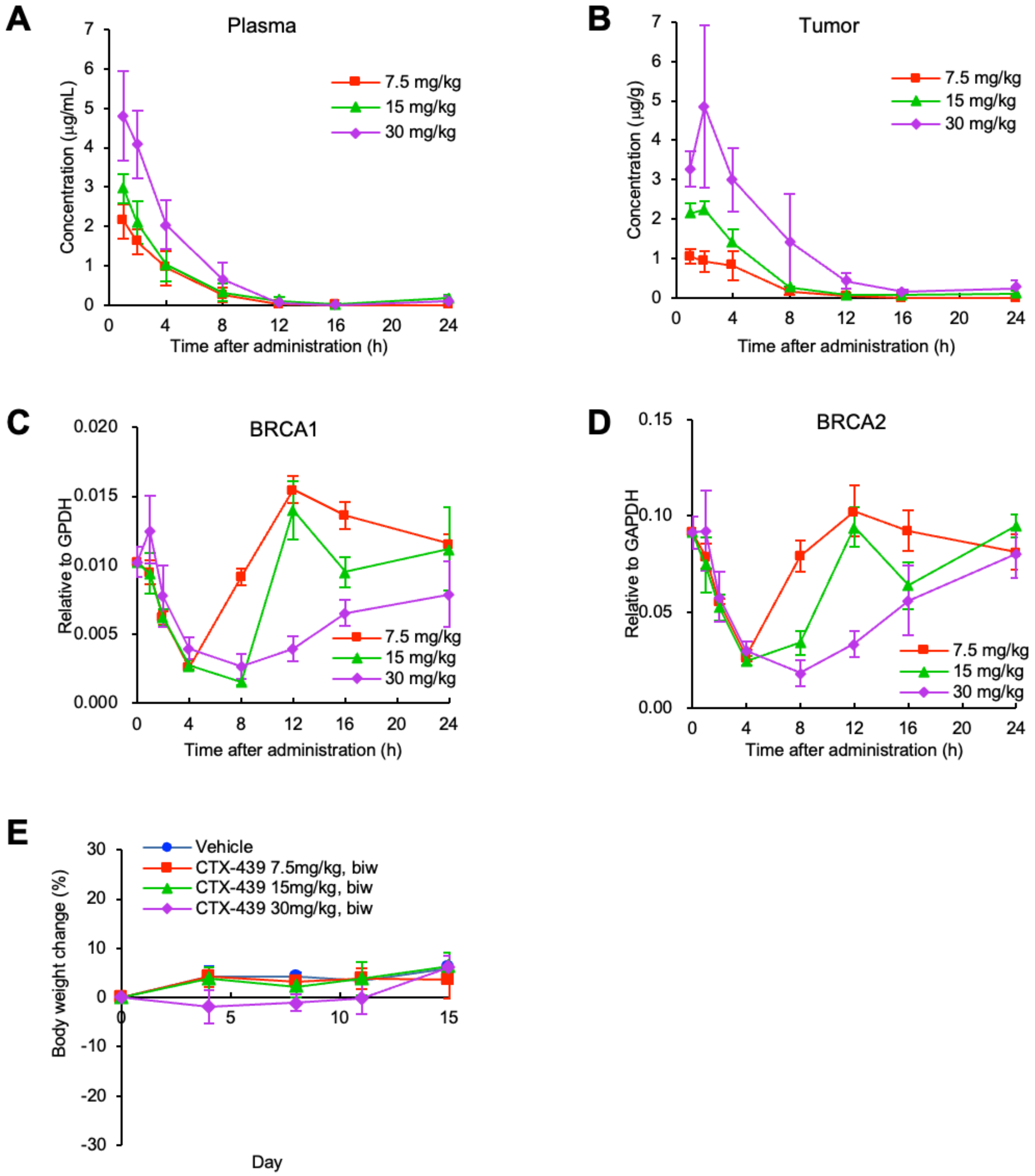


**Figure S3. *In vivo* prolife of CTX-439.**
**A-B**. Plasma (**A**) and intra-tumor (**B**) concentration of CTX-439 at the indicated time after administration at the indicated dose. **C-D**. RT-qPCR analysis of *BRCA1* and *BRCA2* gene in the xenograft tumor cells. Data are shown as mean ± SD (n=3). **E**. Body weight change during CTX-439 *in vivo* treatment (Fig. 1**H**). Data are shown as mean ± SD (n=6).

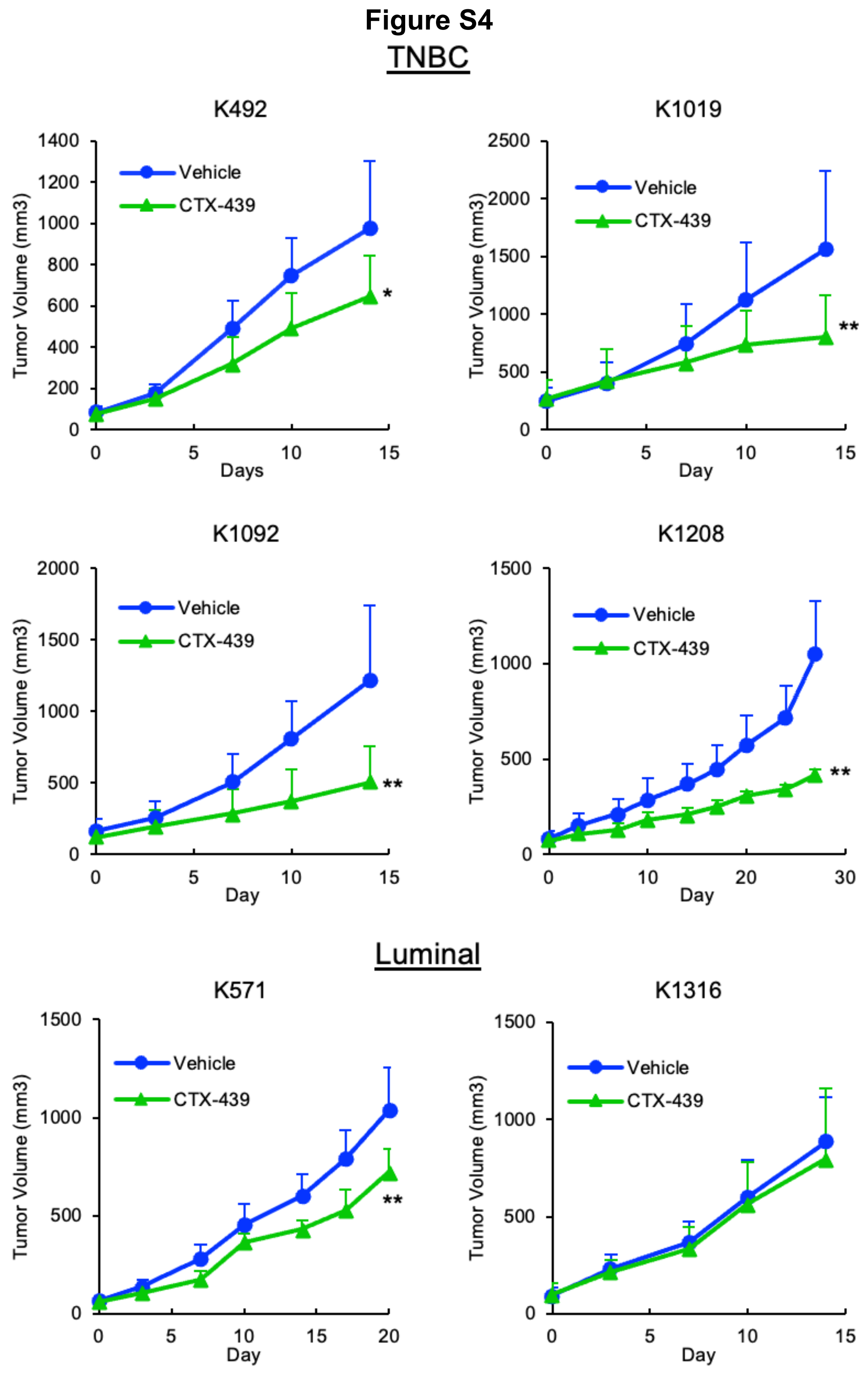


**Figure S4.** ***in vivo*** **efficacy of CTX-439 in breast cancer PDX.**
Breast cancer PDX model mice treated at 15 mg/kg twice weekly. * $p<0.05$, ** $p<0.01$ by Student's *t*-test. Data are shown as mean ± SD (n=4-6).

#### Figure S5

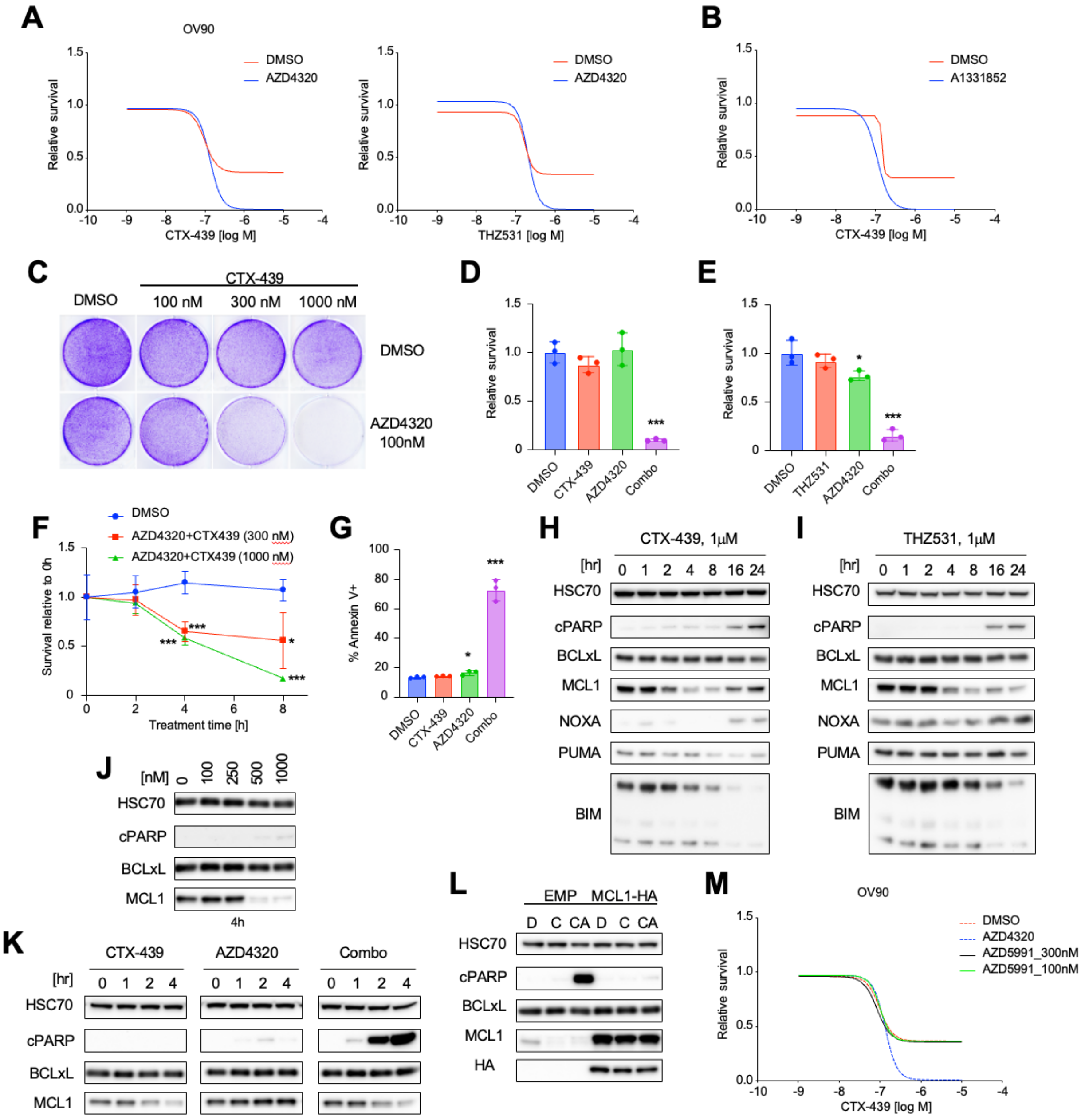


**Figure S5. BCL-2/xL and CDK12/13 dual inhibition induced rapid apoptosis in OV90.**
**A**. Cell survival assay. Cells were treated with serially diluted CTX-439 or THZ531 together with DMSO or AZD4320 for 72h. **B**. Cell survival assay. Cells were treated with serially diluted CTX-439 together with DMSO or A1331852 (30 nM) for 72h. **C**. Crystal violet staining of cells treated in the indicated drug combination for 8h followed by 40h incubation without treatment. **D-E**. Relative cell survival after the indicated drug treatment. Cells were treated with the indicated inhibitors for 8h (**D**) or 10h (**E**) and then surviving cells were counted. **F**. Relative cell counts at the indicated timepoints. **G**. Annexin V staining. Cells were treated with either CTX-439 or AZD4320, or in combination for 6h. **H-I**. Western blot analysis of BCL-2 family proteins. Cells were

treated with CTX-439 (**H**) or THZ531 (**I**) for the indicated time and then lysed. **J-K**. Western blot analysis. Cells were treated with the indicated concentration of CTX-439 for 4 h (**I**) or for the indicated time (**K**). **L**. Western blot analysis of MCL1-over-expressing cells treated with DMSO (D), CTX-439 alone (C) or in combination with AZD4320 (CA). MCL1 was tagged with HA. **M**. Cell survival assay. Cells were treated with serially diluted CTX-439 together with DMSO, AZD4320, or AZD5991 (100 or 300 nM). Unless otherwise specified, CTX-439 at 1 μM and AZD4320 at 100 nM were used. Data are shown as mean ± SD (n=3, **D-G**). * $p$<0.05, ** $p$<0.01, *** $p$<0.001 by Student's *t*-test.

#### Figure S6

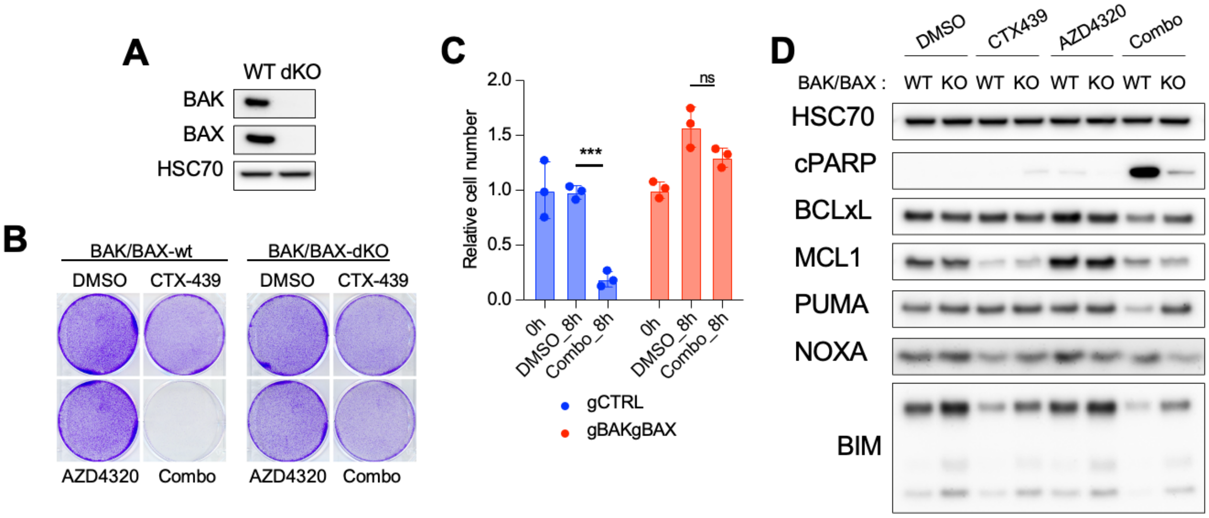


**Figure S6. *BAK/BAX* double-knockout suppressed the rapid induction of apoptosis.**

**A**. Western blot confirmation of *BAK/BAX* double knockout (dKO). **B**. Crystal violet staining. dKO survived even under the combination treatment. **C**. Cell counts 8h post treatment. **D**. Western blot analysis of BCL-2 family protein and cleaved PARP expression in *BAK/BAX*-dKO cells treated with the indicated single or dual agents for 4h. Note that dKO substantially reduced apoptosis induction, while MCL-1 levels were similarly reduced to wt cells. Combination (combo) or single treatment were carried out with CTX-439, 1 $\mu$M and/or AZD4320, 100 nM. Data are shown as mean ± SD (n=3, **C**). *** $p$<0.001 by Student's $t$-test; ns, not significant.

**Figure S7**

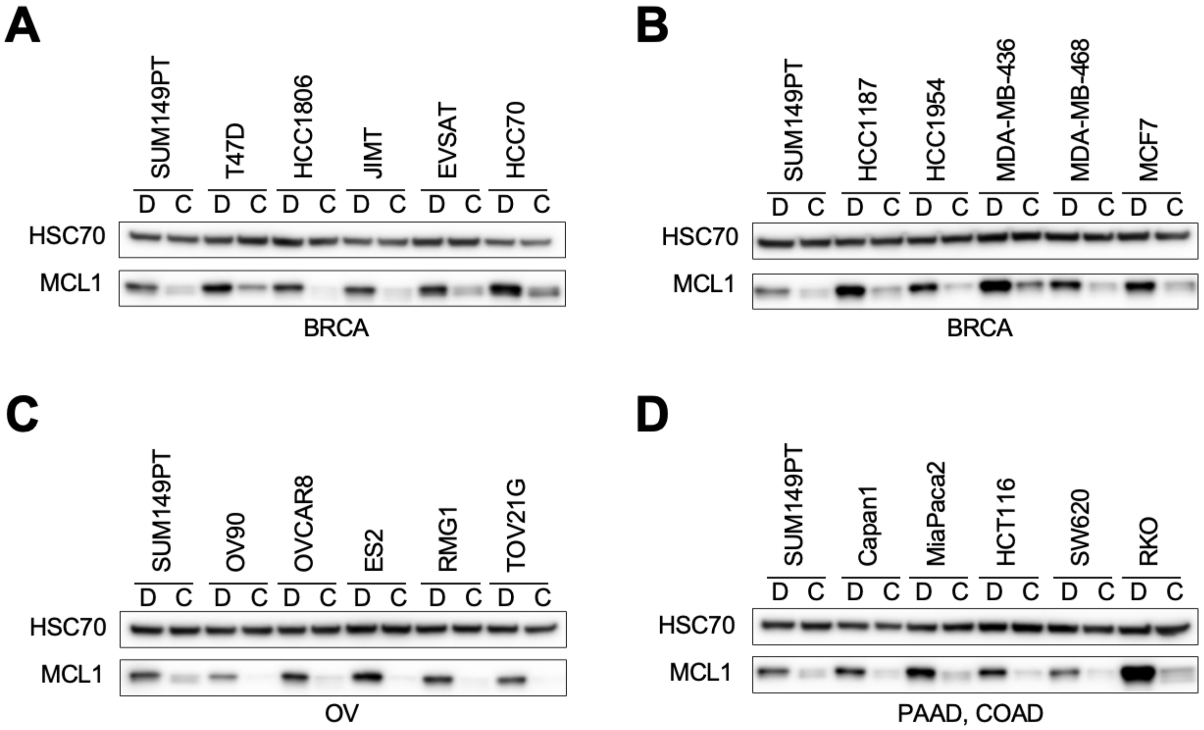


**Figure S7. MCL1 downregulation in a wide range of cancer cell lines.**
**A-C**. Western blot analysis of MCL1 expression in cells treated with CTX-439, 800 nM for 4h. Breast cancer cell lines (**A**-**B**) and ovarian (**C**), pancreatic and colorectal cancer cells lines (**D**) were tested with DMSO (D) or CTX-439 (C). For a reference purpose, the same SUM149PT lysate was loaded to each panel (**A**-**D**).

#### Figure S8

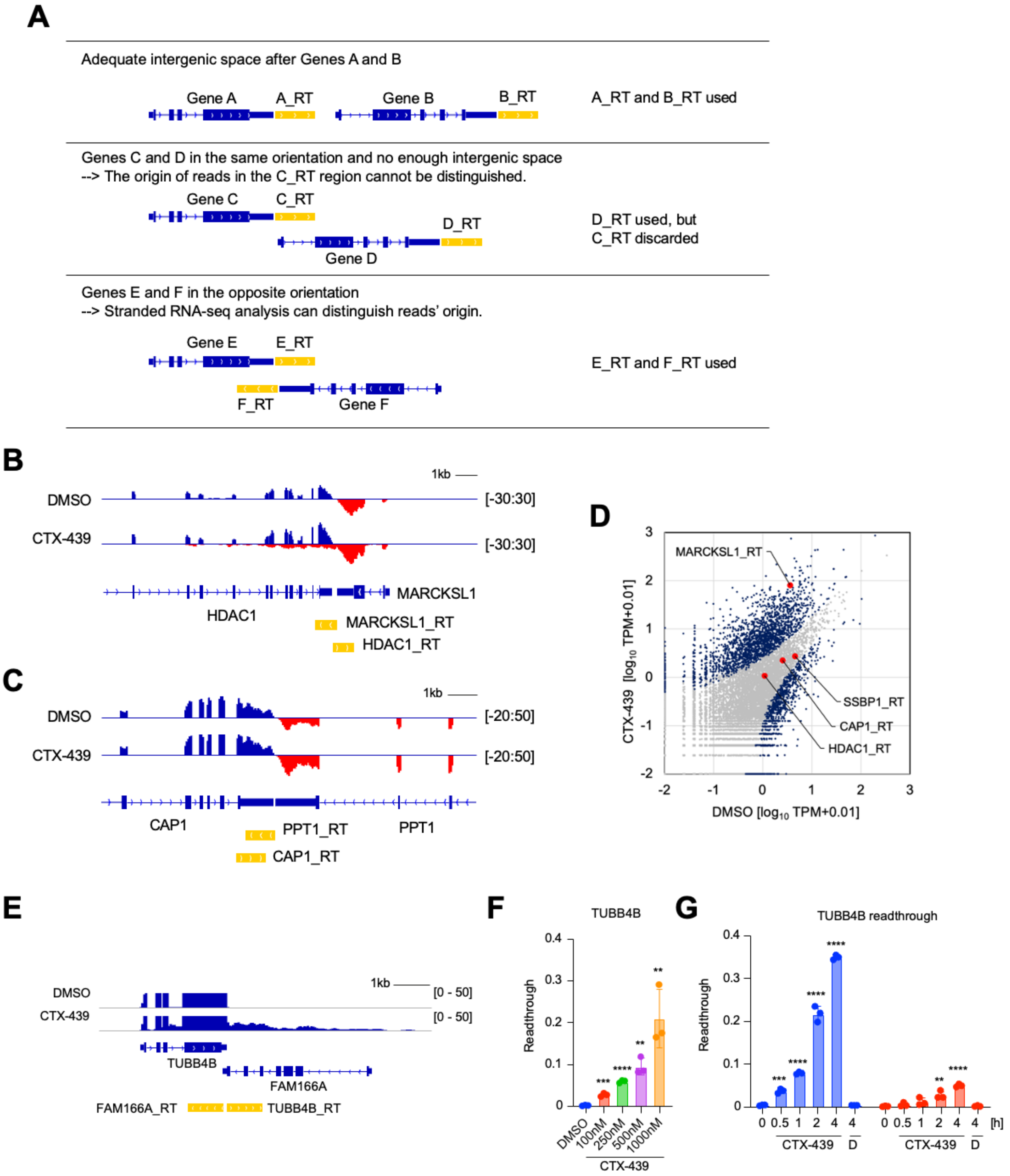


**Figure S8. Defining genomic regions to count readthrough reads by RNA-seq**
**A.** In principle, a 1-kb region immediately after the last exon was set for each protein-coding gene. Following stranded RNA-seq analysis, reads mapped within this region were counted as readthrough transcript. When setting the 1-kb regions, genes were categorized into the following three groups, depending on adjacent gene arrangement. (Top) If there is an adequate intergenic space, a 1-kb region immediately after the last

exon was used. (Middle) If a 1-kb region overlaps with a 3'-adjacent gene that has the same transcriptional orientation, this region was discarded for analysis. This is because read origins cannot be distinguished between the target and the adjacent genes. (Bottom) If a 1-kb region overlaps with a 3'-adjacent gene whose transcriptional orientation is opposite to the target gene, this region was used for analysis. Stranded RNA-seq analysis can distinguish read's origin. **B-C.** Two example loci coding genes in the opposite orientation with a tight intergenic space. The *MARCKSL1* gene (encoded on the - strand) showed readthrough transcription, but the other genes did not. **D**. Scatter plot showing readthrough counts with the example genes in **B** and **C** highlighted. Each dot represents a gene. Blue dots are significantly altered genes. **E**. RNA-seq track showing
*TUBB4B* expression in rRNA-depleted nuclear RNAs. Negative-strand track is omitted. **F-G**. Dose- (**F**) and time-series (**G**) RT-qPCR analysis of *TUBB4B* readthrough transcription. Cells were treated with DMSO (D) or CTX-439 for 4 h at the indicated concentration (**F**) or at 1 μM for the indicated time (**G**). Nuclear and cytoplasmic RNA were then isolated. Data are shown as mean ± SD (n=3, **F-G**). * $p$<0.05, ** $p$<0.01, *** $p$<0.001, **** $p$<0.0001 by Student's *t*-test.

#### Figure S9

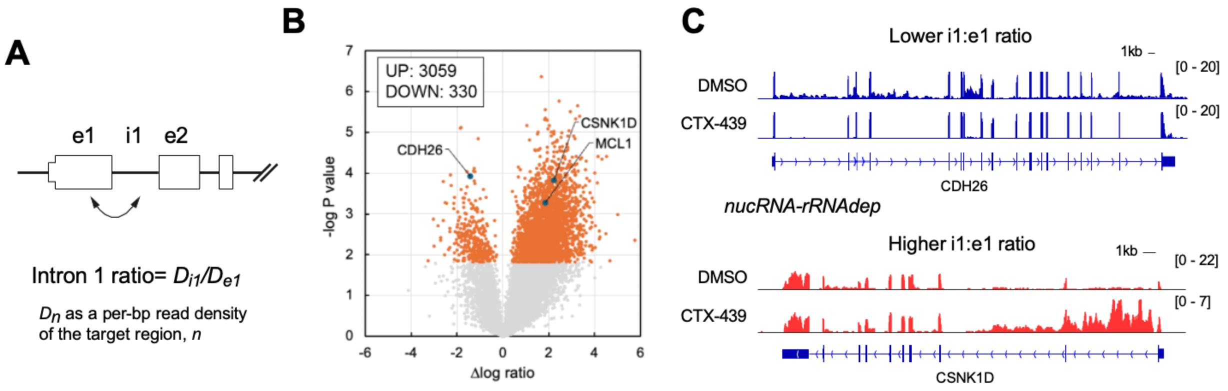


**Figure S9. Intron 1 splicing defect in CTX-439-treated OV90 cells.**
**A**. Schematic of the E1-E2 region. **B.** Volcano plot showing Δlog ratio vs significance. **C**. Gene track of representative genes with reduced (top) and increased (bottom) i1:e1 ratios. Note that signals from + and - strand genes were colored in blue and red, respectively.

**Figure S10**

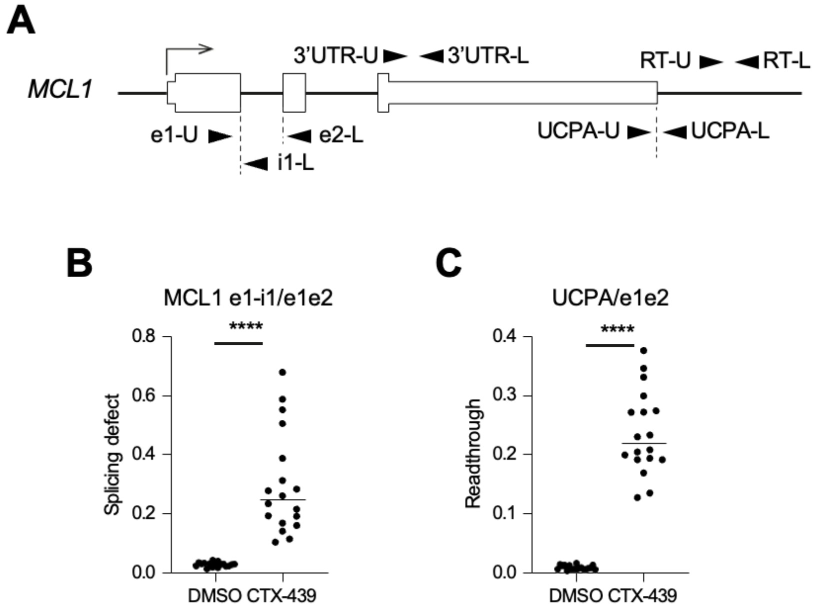


**Figure S10. *MCL1* expression in CTX-439-treated cells.**
**A**. Schematic of the *MCL1* locus. Primers used for quantification were depicted. **B-C**. Unprocessed intron 1 (**B**) and readthrough transcripts (**C**) were quantified using total RNA extracted from cells treated with 800 nM CTX-439 for 4 h. Data were normalized to the processed transcript level (quantified with primers, e1-U and e2-L). 9 breast cancer, 4 ovarian cancer, 3 colon adenocarcinoma, and 2 pancreatic adenocarcinoma cell lines were analyzed. Note that primers, RT-U and RT-L, are used to quantified readthrough, and 3'UTR-U and 3'UTR-L are used for normalization in Figure 3**F**-**G**. **** $p<0.0001$ by Student's *t*-test.

### Figure S11

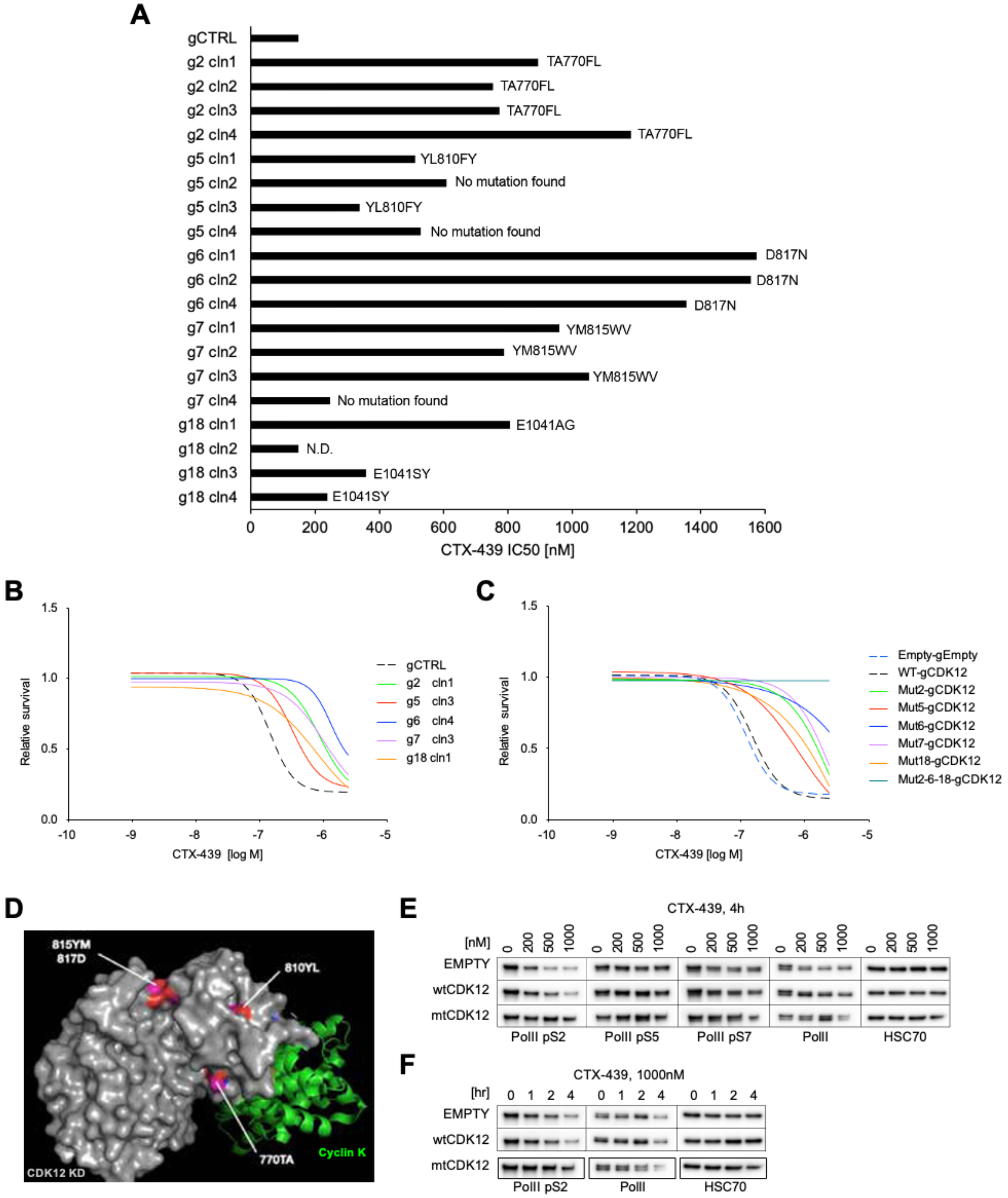


**Figure S11. Identification of CTX-439-resistant CDK12 mutations**

**A**. $IC_{50}$ values and identified coding mutations in the isolated CTX-439-resistant colonies that emerged from gRNA-transduced MDA-MB-436 cells. gRNA and clone IDs were shown. **B**. Cell survival assay of a representative mutant clone from each gRNA. **C**. Cell survival assay. MDA-MB-439 cells were transduced with empty, wildtype or mutant CDK12 cDNA (gRNA-resistant), followed by gCDK12 to knockout the endogenous CDK12. Mut2-6-18 cDNA carries 3 mutations (TA770FL, D817N, E1041AG) and showed high resistance. **D**. The mutations were positioned around the groove of the CDK12 kinase domain. **E-F**. Western blot analysis. Dose (**E**) and time-course (**F**) dependency were examined. Note that mtCDK12-expressgin cells did not reduce the pS2 level.

**Table S1.**
**Kinase inhibitory activity of CTX-439.**

| Kinase | $IC_{50}$ (nM) | Ki (nM) | Km ATP (μM) |
|---|---|---|---|
| CDK12/Cyclin K | 3.1 | 1.4 | 30 |
| CDK13/Cyclin K | 9.2 | 5.5 | 5 |
| CDK2/Cyclin A | 46 | 24 | 20 |
| CDK4/Cyclin D3 | 134 | 69 | 30 |
| CDK6/Cyclin D3 | 551 | 358 | 100 |
| CDK7/Cyclin H | 216 | 105 | 50 |
| CDK9/Cyclin T1 | 60 | 28 | 10 |
| TAOK1 | 4.8 | 2.3 | 30 |
| TAOK3/JIK | 102 | 48 | 50 |
| DYRK1/DYRK1A | 5.5 | 2.6 | 50 |
| DYRK1B | 3.2 | 1.6 | 20 |
| ASK1/MAP3K5 | 14 | 6.5 | 30 |
| JNK2 | 43 | 22 | 20 |

Enzyme inhibitory activities were evaluated using HotSpot™ by Reaction Biology.

**Table S2.**

**$IC_{50}$ values of CTX-439 in cell-based assays using breast and ovarian cancer cell lines.**

| Breast | $IC_{50}$ [nM] | Ovarian | $IC_{50}$ [nM] |
|---|---|---|---|
| AU565 | 21.06 | EFO27 | 51.98 |
| HCC1954 | 44.4 | TYKnu | 53.8 |
| HCC1395 | 46.09 | A2780ADR | 53.98 |
| COLO824 | 49.51 | OVCAR3 | 55.14 |
| MCF7 | 50.49 | A2780 | 57.3 |
| HCC1187 | 57.34 | A2780cis | 59.44 |
| MDAMB468 | 57.88 | TOV112D | 67.74 |
| Hs578T | 65.98 | KURAMOCHI | 72.53 |
| HCC70 | 69.95 | MCAS | 77.86 |
| JIMT1 | 85.54 | TOV21G | 83.2 |
| MDAMB436 | 86.31 | PEO4 | 84.44 |
| MDAMB361 | 88.16 | UWB1 | 86.29 |
| CAL51 | 89.1 | DOV13 | 87.16 |
| CAL851 | 94.37 | OVCAR8 | 91.55 |
| CAMA1 | 95.85 | ES2 | 94.02 |
| HCC38 | 98.53 | JHOS4 | 94.33 |
| SUM149PT | 100.5 | OC314 | 109.6 |
| MDAMB453 | 100.9 | JHOS2 | 109.6 |
| MFM223 | 105.1 | OV90 | 116 |
| OCUBM | 109.9 | OVCAR5 | 127 |
| HCC1806 | 111 | EFO21 | 130.8 |
| MDAMB231 | 118.3 | OVMIU | 136.2 |
| HCC1428 | 124.1 | OVISE | 155.7 |
| HCC1937 | 142.9 | OVCAR4 | 162.1 |
| MDAMB415 | 144.6 | RMGI | 162.4 |
| T47D | 162.3 | Caov4 | 175.1 |
| EVSAT | 236.5 | Hey | 190.9 |
| ZR751 | 285 | OAW42 | 230.8 |

**Table S3.**
**PDX (CrownBio) characteristics**

| PDX ID | Subject information | | | | | | | Xenograft information | | | |
|---|---|---|---|---|---|---|---|---|---|---|---|
| | Diagnosis | NAC | Grade | ER | PR | HER2 | Metastasis | ER | PR | HER2 | Ki-67 |
| BR9457 | IDC | - | 3A | Low | Low | Neg | Lung | Neg | Neg | 1+ | 40 |
| BR9465 | IDC | + | 3C | Low | Low | Neg | Lung | Neg | Neg | Neg | 90 |

These models are considered as TNBC based on the reference: Acs, Balazs, et al. The Lancet Regional Health-Europe 40 (2024).

**Table S4.**

**PDX (Kyoto) characteristics**

| PDX ID | NAC | Grade | ER (%) | PR (%) | HER2 (score) | Ki-67 (%) |
|---|---|---|---|---|---|---|
| K492 * | + (Grade 1b) | 3 | Allred score 3 | 0 | 0 | 45 |
| K571 | + | 3 | 30 | 0 | 0 | 69 |
| K1019 | + (Grade 0) | 3 | 0 | 0 | 0 | 69 |
| K1092 | + (Grade 0) | 3 | 0 | 0 | 1+ | 76 |
| K1208 | + | 3 | 0 | 0 | 2+ | 43 |
| K1316 | + | 3 | 3 | 0 | 3+ | 71 |

* K492 with Allred score 3 (PS 1 + IS 2), corresponding to < 1% ER expression, were considered ER-negative and classified as TNBC in accordance with ASCO/CAP guidelines.

**Table S5.**
**gRNA sequence**

| | Identifier | Sequence (5' - 3') |
|---|---|---|
| 1 | CDK12-e4-1147297119 | GGCCAAGTATATAAAGCCA |
| 2 | CDK12-e5-1147297333 | AGACTAGACAATGAGAAAG |
| 3 | CDK12-e5-1147297336 | TTTCACGAATGGCTGTGAT |
| 4 | CDK12-e5-1147297337 | AGGATTTTGATTTCACGAA |
| 5 | CDK12-e5-1147297343 | GATGCACTGGATTTCAAGA |
| 6 | CDK12-e6-1147298140 | TACTCAAATACAAGGTAAA |
| 7 | CDK12-e6-1147298141 | TACCTTGTATTTGAGTATA |
| 8 | CDK12-e6-1147298142 | GTCCATATACTCAAATACA |
| 9 | CDK12-e6-1147298145 | TAGCAGTCCCATTAAGTCA |
| 10 | CDK12-e6-1147298146 | TAATGGGACTGCTAGAATC |
| 11 | CDK12-e6-1147298147 | GGACTGCTAGAATCTGGTT |
| 12 | CDK12-e6-1147298149 | TTTCATGAACGACTTGATA |
| 13 | CDK12-e6-1147298151 | TCATGAAACAGCTAATGGA |
| 14 | CDK12-e6-1147299314 | TAGCAGTAGTTCTGGAGGT |
| 15 | CDK12-e7-1147299078 | AAATCAAACTAGCAGATTT |
| 16 | CDK12-e8-1147299321 | CCATAGATGTTTGGAGCTG |
| 17 | CDK12-e8-1147299825 | AGGCCACACAGCTGGACAA |
| 18 | CDK12-e8-1147299827 | TAACATCAGGCCACACAGC |
| 19 | CDK12-e12-1147300722 | TCCTGCCAGTGGGGGAGGC |
| 20 | CDK12-e12-1147300723 | GCAATCCTGCCAGTGGGGG |
| 21 | CDK12-e12-1147300724 | ATGGCAATCCTGCCAGTGG |
| 22 | CDK12-e12-1147300725 | CATGGCAATCCTGCCAGTG |
| 23 | CDK12-e12-1147300726 | TCATGGCAATCCTGCCAGT |
| 24 | CDK12-e12-1147300728 | GCAGGATTGCCATGAGTTG |
| 25 | CDK12-e12-1147300731 | GGCGACGTCAGCGACAAAG |
| 26 | CDK12-e12-1147300740 | AGCAGGCTGAGGTGGTGCT |
| 27 | CDK12-e12-1147300746 | CAGCCTGCTCCTGGCAAGG |
| 28 | CDK12-e12-1147300748 | CTCCTGGCAAGGTGGAGTC |
| 29 | CDK12-e12-1147300755 | CTGGGGCTGGGGATGCAAT |
| | gCDK12_v3_6-2 | GTTCACGCAGTCGTCATTC |
| | gCDK12_v3_6-4 | TGAGGCTGCCCCAAAAGGG |

**Table S6.**

**PCR primers for plasmid construction and MCL1 genotyping**

| Target | Forward primer (5' - 3') | Reverse primer (5' - 3') |
|---|---|---|
| MCL1 cloning | AGGTGTCGTGGGAAGCTTggCCACCATGTTTGGCCTCAAAAGAAA | CCGATCTTGTTGCGTCAggCTAAGCGTAATCTGGTACGTCGTATGGGTATCCTCCGGAcCCGCCGCCTCTTATTAGATATGCCAAAC |
| CDK12 cloning | AGGTGTCGTGGGAAGCTTggccaccATGTACCCATACGACGTACCAGATTACGCTCCCAATTCAGAGAGACATGG | CCGATCTTGTTGCGTCAggCTAGTAAGGAACTCCTCTCCC |
| Calabrese transfer to pKLV2 | TTGGTTAGTACCGGGCCCTAGAGGGCCTATTTCCCATGAT | GGGCGTTTGATCGTTTGCCGAAAAAAAGCACCGACTCGGTGCCAC |
| MCL1-purobpA | GGGGCAGGATTGTGACTCT | CCAATCAGAGCCCATTATTTGT |

**Table S7.**
**qPCR primer sequences**

| Target gene | Forward primer (5' - 3') | Reverse primer (5' - 3') |
|---|---|---|
| GAPDH * | TGTAGTTGAGGTCAATGAAGGG | ACATCGCTCAGACACCATG |
| BRCA1 | GCTCTTCGCGTTGAAGAAGTA | CACTGTGAAGGCCCTTTCTT |
| BRCA2 | CGCTGCAACAAAGCAGATTTA | TTCAGATTCTTCTGCAGGTTCA |
| MCL1_3' UTR | CCTCCATAGCTTCCCAAACA | AAAAGCAAGTGGCAAGAGGA |
| MCL1_RT | TCCATGGTGATTCAGCTCAC | TGAAGTTCTTGGCAGCAGTG |
| MCL1_UCPA | GCTTTTGCTTTGAATTCACCT | TGTTGGTTGTCAAATGTAGTGTG |
| MCL1_E1-I1 | CTTACGACGGGTTGGGGAT | AGTGAGGCCTTGGCGATTAA |
| MCL1_E1-E2 | CTTACGACGGGTTGGGGAT | GGATCATCACTCGAGACAACG |
| MCL1 | ATGTCCAGTTTCCGAAGCAT | CCAAGGACACAAAGCCAATG |
| TUBB4B | GCAGCTGGAACGCATCAAC | GGGCTCCAGATCCACGAG |
| TUBB4B_RT | CGAATGTCAGGGCGTAGTTG | CTTCTCTGAAGCCTGGGTGT |
| GAPDH ** | CGCTTCGCTCTCTGCTCCTCCTGT | GGTGACCAGGCGCCCAATACGA |

* Used in Fig. 1 and Supplementary Fig. 3.
** Used in Figs. 3, 4 and 6.

**Table S8.**
**Antibody used in this study**

| Antibodies | Supplier | Identifier |
|---|---|---|
| pS423-CDK12 | Chordia Therapeutics Inc. | S87A14I-85 |
| CDK12 | Proteintech | 26816-1-AP |
| pS2 polII CTD | MERCK | 04-1571 |
| pS5 polII CTD | MERCK | 04-1572 |
| pS7 polII CTD | MERCK | 04-1570 |
| polII CTD | Convance | MMS-126R-500 |
| β-actin | Cell Signaling Technology | #5125 |
| BRCA1 | Cell Signaling Technology | #9010 |
| BRCA2 | Cell Signaling Technology | #10741 |
| RAD51 | Cell Signaling Technology | #8875 |
| γH2AX | Cell Signaling Technology | #2577 |
| Cleaved Caspase-3 | Cell Signaling Technology | #9661 |
| MCL1 * | BD Pharmingen | #559027 |
| HSC70 | Santa Cruz | sc-7298 |
| Cleaved PARP | Cell Signaling Technology | #5625 |
| BCL-xL | Cell Signaling Technology | #2764 |
| MCL1 ** | Cell Signaling Technology | #94296 |
| NOXA | Cell Signaling Technology | #14766 |
| PUMA | Cell Signaling Technology | #98672 |
| BIM | Cell Signaling Technology | #2933 |
| HA | BioLegend | 901502 |
| BAK | Cell Signaling Technology | #12105 |
| BAX | Cell Signaling Technology | #5023 |
| T2A | MERCK | MABE1923 |
| CDK12 | Cell Signaling Technology | 11973 |
| PolII pS2 | Cell Signaling Technology | #13499 |

| | | |
|---|---|---|
| PolII pS5 | Cell Signaling Technology | #13523 |
| PolII pS7 | Cell Signaling Technology | #13780 |
| PolII | Cell Signaling Technology | #14958 |
| anti-Rabbit IgG, HRP | Cell Signaling Technology | #7074 |
| anti-Mouse IgG, HRP | Cell Signaling Technology | #7076 |

* Used in Fig. 7.
** Used in Figs. 2, 4 and 6, and Supplementary Figs. 4, 5 and 6

**Data S1. (separate file)**
Kinase affinity panel of CTX-439 against 468 kinases

**Data S2. (separate file)**
DESeq2_result_CTX-439_100nM

**Data S3. (separate file)**
SUM149PT_CRISPRa.gene_summary
